# X-Cell: Scaling Causal Perturbation Prediction Across Diverse Cellular Contexts via Diffusion Language Models

**DOI:** 10.64898/2026.03.18.712807

**Authors:** Chloe Wang, Mehran Karimzadeh, Neal G. Ravindra, Lexi R. Bounds, Nader Alerasool, Ann C. Huang, Shihao Ma, Daniel R. Gulbranson, Haotian Cui, Yongju Lee, Anusuya Arjavalingam, Elliot J. MacKrell, Matthew S. Wilken, Jieming Chen, Benjamin W. Herken, Jesse A. Weber, Massimo M. Onesto, Barbara Gonzalez-Teran, Nicole F. Leung, Sally Yu Shi, Byron J. Smith, Sharon K. Lam, Adam Barner, Philip Wright, Elizabeth M. Rumsey, Soohong Kim, Rene V. Sit, Adam J. Litterman, Ci Chu, Bo Wang

## Abstract

Causal models of cellular systems hold the promise to empower broad biological discovery, including the systematic identification of novel targets for drug discovery. Predicting how genetic and pathway perturbations reshape gene expression across diverse cellular contexts is a prerequisite for building generalizable cellular foundation models. However, current methods typically fail to extrapolate beyond their training distributions because they rely predominantly on observational expression atlases rather than interventional perturbation data. We present X-Atlas/Pisces, the largest genome-wide CRISPRi Perturb-seq compendium to date, comprising 25.6 million perturbed single-cell transcriptomes across 16 biologically diverse contexts, including widely used cell lines, induced pluripotent stem cells (iPSCs), resting and CD3/CD28 activated Jurkat T lymphoma cells, and multi-lineage differentiating iPSCs. Leveraging this resource, we develop X-Cell, a diffusion language model that predicts perturbation responses by iteratively refining control-to-perturbed state transitions through cross-attention to multi-modal biological priors derived from natural language, protein language models, interaction networks, genetic dependency maps, and morphological profiles. X-Cell outperforms existing state-of-the-art models by up to five-fold on key metrics such as Pearson **Δ** (correlation between predicted and observed perturbation-induced log-fold changes), and demonstrates zero-shot prediction of T cell inactivating perturbations in stimulated Jurkat cells. We scale X-Cell to 4.9 billion parameters (X-Cell-Ultra), the largest causal perturbation model to date. We demonstrate for the first time that perturbation prediction follows power-law scaling with an exponent matching large language models. X-Cell-Ultra demonstrates zero-shot generalization to novel biological contexts, including unseen iPSC-derived melanocyte progenitors and primary human CD4^+^ T cells from multiple donors, and outperforms all baselines after self-supervised test-time adaptation. These results demonstrate that coordinated scaling of causal perturbation data and model capacity yields foundation models capable of generalizable perturbation prediction across cellular contexts, with potential applications for improving computational target identification, validation, and context-specific therapeutic prioritization.

## 1 Introduction

A central challenge in modern drug discovery is anticipating how cellular systems will respond to genetic or chemical interventions. Accurately predicting the effects of these cellular perturbations has direct translational value: it enables the computational identification and prioritization of therapeutic targets, the validation of gene-disease links before experimental follow-up, and the matching of interventions to patients most likely to benefit from a specific therapeutic mechanism [1]. However, modeling these responses is fundamentally complex because cells are governed by deeply layered regulatory systems spanning chromatin architecture [2], transcriptional control [3], RNA processing [4], post-translational regulation [5], and disease-associated molecular states [6, 7]. To navigate this biological complexity, recent advances in high-content perturbation screens and machine learning have motivated the development of foundation models that aim to capture these interconnected regulatory layers. Critically, most existing single-cell foundation models are trained on observational transcriptomic atlases that capture correlative co-expression patterns, conflating statistical association with causation [8–10]. Genome-wide perturbation screens, by contrast, provide interventional ground truth by directly measuring the causal consequences of genetic perturbations and resolving directed regulatory dependencies [11–13]. Models trained on such causal data can learn the mechanistic structure required for reliable out-of-distribution generalization tasks and predict how untested perturbations reshape transcriptional programs in novel cellular contexts [14].

Here, we consider a perturbation foundation model as one that can accurately predict genome-scale transcriptional responses to perturbations of any gene or pathway in a given cellular context, including contexts not present in training data. Such predictive models are essential because the combinatorial space of possible perturbations — including across genes, cell types, and chemical conditions — is far too large to explore experimentally. Early efforts in perturbation prediction relied on probabilistic latent variable models to characterize the basal cellular state and represent structured shifts in gene expression space conditioned on perturbations [15–17]. Subsequent approaches incorporated prior knowledge of gene–gene interactions through graph-based modeling [18], as well as broader biological knowledge encoded in textual gene descriptions combined with attention-based architectures [19–21]. In parallel, advances leveraging large single-cell foundation models [8–10, 22–24], along with optimal transport and ODE-based frameworks for modeling perturbation dynamics at the single-cell level [25–29], have continued to push the boundaries of predictive performance. The field has witnessed both substantial methodological advances and limitations. More expressive models often achieve strong predictive performance [30]. However, evidence also suggests that these models do not reliably surpass simple baselines such as mean-perturbation of the same context [31, 32]. While many models perform well in previously observed settings (e.g., known cell types or perturbations), performance gaps remain when generalizing to unseen perturbations or cell types not encountered during training [33].

Recent efforts have focused on scaling foundation models and language-based perturbation models but have not consistently yielded meaningful improvements over smaller counterparts [22, 23, 34]. This suggests that increasing model parameters alone may be insufficient to close the generalization gap. Cross-context generalization is challenging because gene-regulatory causality is *inherently* contextdependent, varying widely across tissues and cell types [35]. Therefore, building a model that captures these structured, context-dependent regulatory dependencies requires the coordinated scaling of both model capacity and *causal* interventional data of unprecedented scale [36, 37]. To this end, we present the X-Atlas/Pisces dataset, the largest genome-wide CRISPRi single-cell perturbation compendium generated to date, comprised of 25.6 million perturbed cells. X-Atlas/Pisces expands beyond single-context guideposts and integrates seven genome-scale Perturb-seq screens across highly diverse biological states, capturing transcriptome-wide perturbation effects including changes in regulatory networks governing metabolism, immune signaling, and dynamic multi-lineage differentiation trajectories. By mapping how biological network dependencies rewire across these diverse environments, X-Atlas/Pisces establishes the necessary structural foundation for context-aware predictions.

Leveraging this compendium, we introduce X-Cell, a diffusion language model (LM) capturing the transcriptomic shift from a control state to a perturbed state via an iterative diffusion process incorporating multi-modal biological priors directly into the generative architecture through cross-attention [19, 38]. We demonstrate that X-Cell consistently outperforms standard predictive baselines and achieves state-of-theart performance on held-out data. Further, the architecture exhibits strong scaling behavior, with model performance consistently improving alongside increased training data and capacity, and demonstrating that X-Cell-Ultra can effectively exploit the increasing number of single-cell perturbation datasets. This scale unlocks state-of-the-art zero-shot generalization, enabling accurate prediction of single-cell transcriptomes in entirely unseen cell types and primary human cells.

## 2 Results

### 2.1 Generation of the X-Atlas/Pisces dataset and X-Cell model

To capture context-dependent perturbation effects, we generated X-Atlas/Pisces, the largest genomewide CRISPRi Perturb-seq resource to date. Expanding our initial X-Atlas/Orion dataset (HCT116 and HEK293T cell lines) [39], Pisces maps perturbations across 25.6 million single-cell transcriptomes from induced pluripotent stem cells (iPSCs), hepatocellular carcinoma cells (HepG2), T lymphoblastic leukemia cells (Jurkat) in resting and CD3/CD28-stimulated states, and multi-lineage differentiating iPSCs (iPSC Multi-Diff) (Figure 1A). Four screens utilized an optimized FiCS Perturb-seq protocol enabling FACS after cryopreservation, adapted to yield high quality data for fragile cell types like HepG2 and Jurkat (Section 4.2.7). For the iPSC Multi-Diff screen, we utilized probe-based Flex Perturb-seq. Flex Perturb-seq provides formaldehyde compatibility, preserving fragile and diverse cell types within a single experiment, and enables FACS-based enrichment of desired cell populations post-fixation. Additionally, sample barcoding enables super-loading; we executed our largest single screen using only 20–25% of standard 10X lanes while achieving *>*50% dual-guide cell recovery yield (Table 1). The iPSC Multi-Diff screen contained 10 annotated cell types (Supplementary Figure 1) reflecting early differentiating lineages after exit from pluripotency. In total, X-Atlas/Pisces contains genome-scale perturbations in 16 cellular contexts (cell type and exposure), achieving *>*152,000 unique conditions for ML training and evaluation.

**Table 1.** Summary of screens in X-Atlas/Pisces. Single-cell assay and # GEMs are related to single-cell library preparation. # aligned cells (total), Mean reads per cell, Median UMIs per cell, and Median genes per cell are related to data sensitivity. Mean CRISPR reads per cell, Median CRISPR UMIs per cell, # cells with two sgRNAs (expected pair), % cells with two sgRNAs (expected pair), and Median number cells per perturbation are related to efficiency of sgRNA capture. Median on-target KD % per perturbation is the primary readout of CRISPRi efficiency. % of perturbations with significant phenotype is related to quantification of secondary effects from perturbation. For details on each metric, see Section 4.2.10.

| Metric | HCT116 | HEK293T | HepG2 | iPSC |
| --- | --- | --- | --- | --- |
| Single-cell assay | GEM-X 5' | GEM-X 5' | GEM-X 5' | GEM-X 5' |
| # GEMs | 109 | 223 | 167 | 199 |
| # cells with two sgRNAs (expected pair) | 3,409,169 | 4,534,299 | 2,642,794 | 4,224,716 |
| # aligned cells (total) | 9,886,825 | 21,550,681 | 8,278,581 | 12,629,592 |
| % cells with two sgRNAs (expected pair) | 34.48 | 21.04 | 31.92 | 33.45 |
| Mean reads per cell | 54,826 | 63,426 | 94,132 | 111,044 |
| Median UMIs per cell | 19,416 | 16,557 | 50,798 | 33,291 |
| Median genes per cell | 5,387 | 5,871 | 6,708 | 7,525 |
| Mean CRISPR reads per cell | 7,610 | 7,263 | 6,708 | 13,449 |
| Median CRISPR UMIs per cell | 950 | 184 | 1,944 | 1,522 |
| # perturbations in screen | 18,924 | 18,312 | 9,735 | 10,095 |
| Median number cells per perturbation | 150 | 200 | 225 | 353 |
| Median on-target KD % per perturbation | 69.95 | 48.10 | 85.11 | 82.18 |
| % of perturbations with significant phenotype | 31.41 | 25.32 | 60.50 | 72.80 |

| Metric | Jurkat Resting | Jurkat Active | iPSC Multi-Diff |
| --- | --- | --- | --- |
| Single-cell assay | GEM-X 5' | GEM-X 5' | Flex |
| # GEMs | 139 | 145 | 34 |
| # cells with two sgRNAs (expected pair) | 2,837,716 | 2,795,555 | 5,125,032 |
| # aligned cells (total) | 8,730,868 | 9,706,890 | 9,978,102 |
| % cells with two sgRNAs (expected pair) | 32.5 | 28.8 | 51.36 |
| Mean reads per cell | 87,882 | 81,645 | 49,089 |
| Median UMIs per cell | 31,896 | 25,478 | 21,239 |
| Median genes per cell | 6,872 | 6,521 | 7,096 |
| Mean CRISPR reads per cell | 9,256 | 8,454 | 5,029 |
| Median CRISPR UMIs per cell | 911 | 492 | 665 |
| # perturbations in screen | 10,872 | 10,878 | 12,175 |
| Median number cells per perturbation | 215 | 210 | 287 |
| Median on-target KD % per perturbation | 78.69 | 70.60 | 95.68 |
| % of perturbations with significant phenotype | 59.12 | 51.73 | 60.33 |

**Fig. 1.**
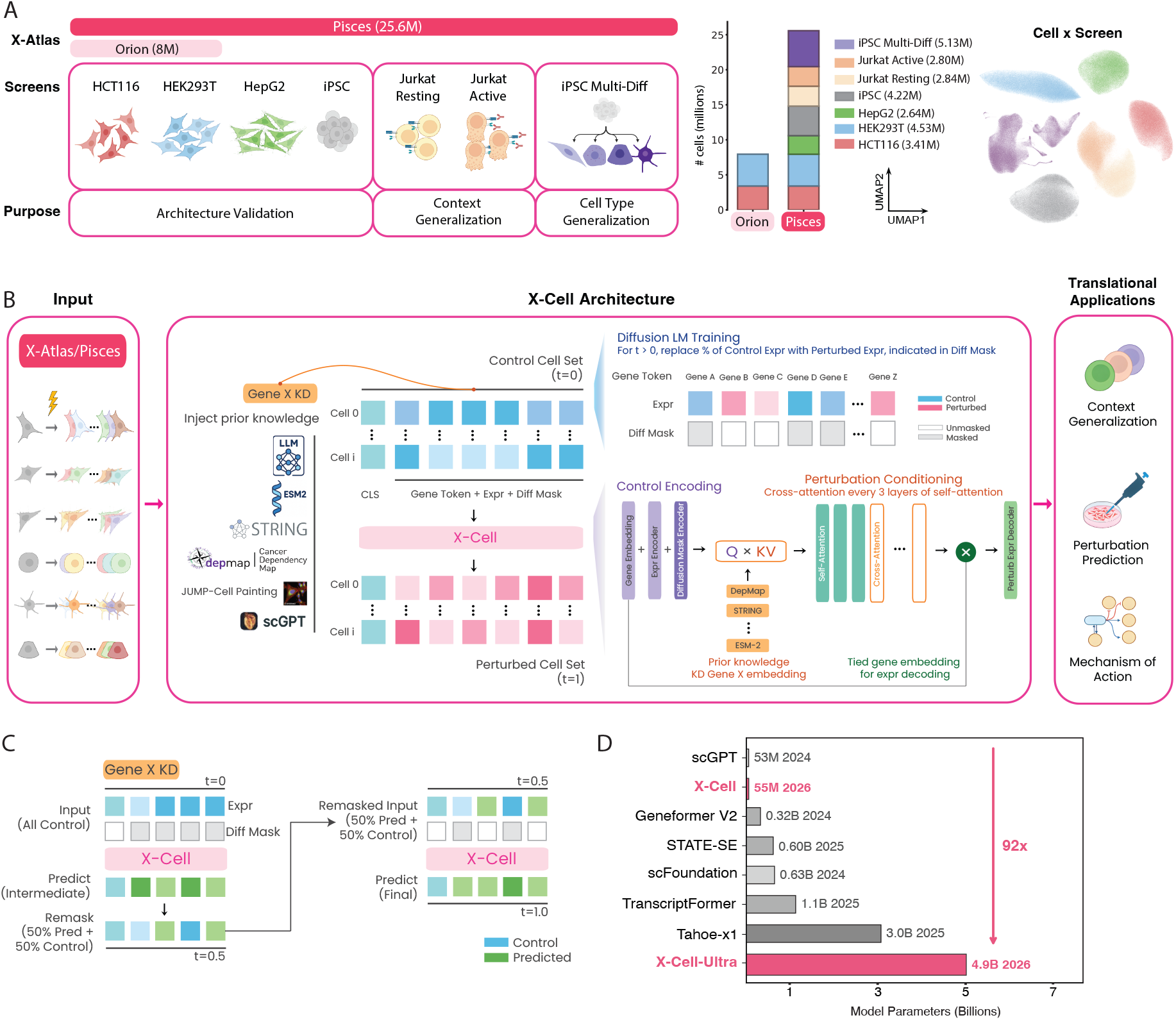
Overview of X-Atlas/Pisces and X-Cell. **(A)** Schematic of X-Atlas/Pisces, which contains seven genome-scale CRISPRi Perturb-seq screens in HCT116, HEK293T, HepG2, iPSC, Jurkat Resting, Jurkat Active, and iPSC Multi-Differentiation (iPSC Multi-Diff) (left). Size comparison of X-Atlas/Orion (8M cells) versus X-Atlas/Pisces (25.6M cells) (middle). UMAP of cells in X-Atlas/Pisces, colored by screen. For visualization, the dataset was downsampled to 1M total cells while preserving the relative proportions of each original screen. Within this subset, 5% of the cells are non-targeting controls and the remainder are perturbed cells. **(B)** X-Cell combines diffusion LM training and cross-attention to prior knowledge to predict perturbed cell states from control cell sets. Six prior knowledge sources include pre-trained embeddings from LLM [19], ESM-2 [38], STRING [40, 41], DepMap [42], JUMP-Cell Painting [43], and scGPT [8] (Section 4.1.2). The main architecture consists of stacked self-attention blocks that encode control cell sets, with interleaved cross-attention to prior knowledge embeddings. X-Cell enables diffusion-style training by randomly replacing 25%, 50%, or 75% of control gene expression values with ground-truth perturbed values, and providing a binary Diff Mask to indicate the revealed positions (Section 4.1.3). **(C)** During inference, X-Cell refines predictions via iterative diffusion by remasking part of its output as input for subsequent generation steps. **(D)** X-Cell scales from 55M parameters (X-Cell) to 4.9B parameters (X-Cell-Ultra), exceeding the size of existing single-cell foundation models.

The large data volume and diversity of X-Atlas/Pisces enabled us to develop X-Cell, a diffusion language model pre-trained across diverse cell lines and perturbations in Pisces (Figure 1B). Because perturbational cascades are constrained by underlying regulatory networks, we designed X-Cell to incorporate multi-modal biological priors directly into the generative process through cross-attention (Figure 1B). These priors are encoded as pre-trained embeddings derived from large language models (GenePT) [19], protein language models (ESM-2) [38], interaction networks (STRING) [40, 41], cancer dependency maps (DepMap) [42], morphological profiling data (JUMP-Cell Painting) [43], and single-cell foundation models [8]. Furthermore, X-Cell models the transcriptomic shift from a control state to a perturbed state via an iterative diffusion process. Predictions are progressively refined across intermediate steps through remasking (Figure 1C). To examine scaling effects, we trained two model variants: X-Cell and X-Cell-Ultra. X-Cell-Ultra contains 4.9 billion parameters, enabling perturbation modeling at unprecedented scale (Figure 1D). To systematically evaluate performance, we partitioned X-Atlas/Pisces into three task-specific regimes: (1) stable cell lines for architecture validation, (2) Jurkat resting and stimulated states for context generalization, and (3) the iPSC Multi-Diff dataset for zero-shot cell type generalization (Section 4.1.7).

### 2.2 X-Atlas captures conserved and context-dependent regulatory networks

The screens in X-Atlas/Pisces yielded high data quality and robust biological signal. Across all seven screens, we achieved deep transcriptomic coverage (median 25,478 UMIs and 6,708 genes per cell) and successful target knockdown (median 78.7%) (Figure 2A). We assessed whether integrating diverse biological states improves the recovery of known gene regulatory networks. Combining perturbation data from one to seven screens yielded a strict, monotonic increase in the recall of established CORUM protein complexes and STRING protein-protein interactions (Figure 2B). Furthermore, for a given number of cells, sampling across multiple contexts improved recall more than deeply sampling a single context (Supplementary Figure 2A).

**Fig. 2.**
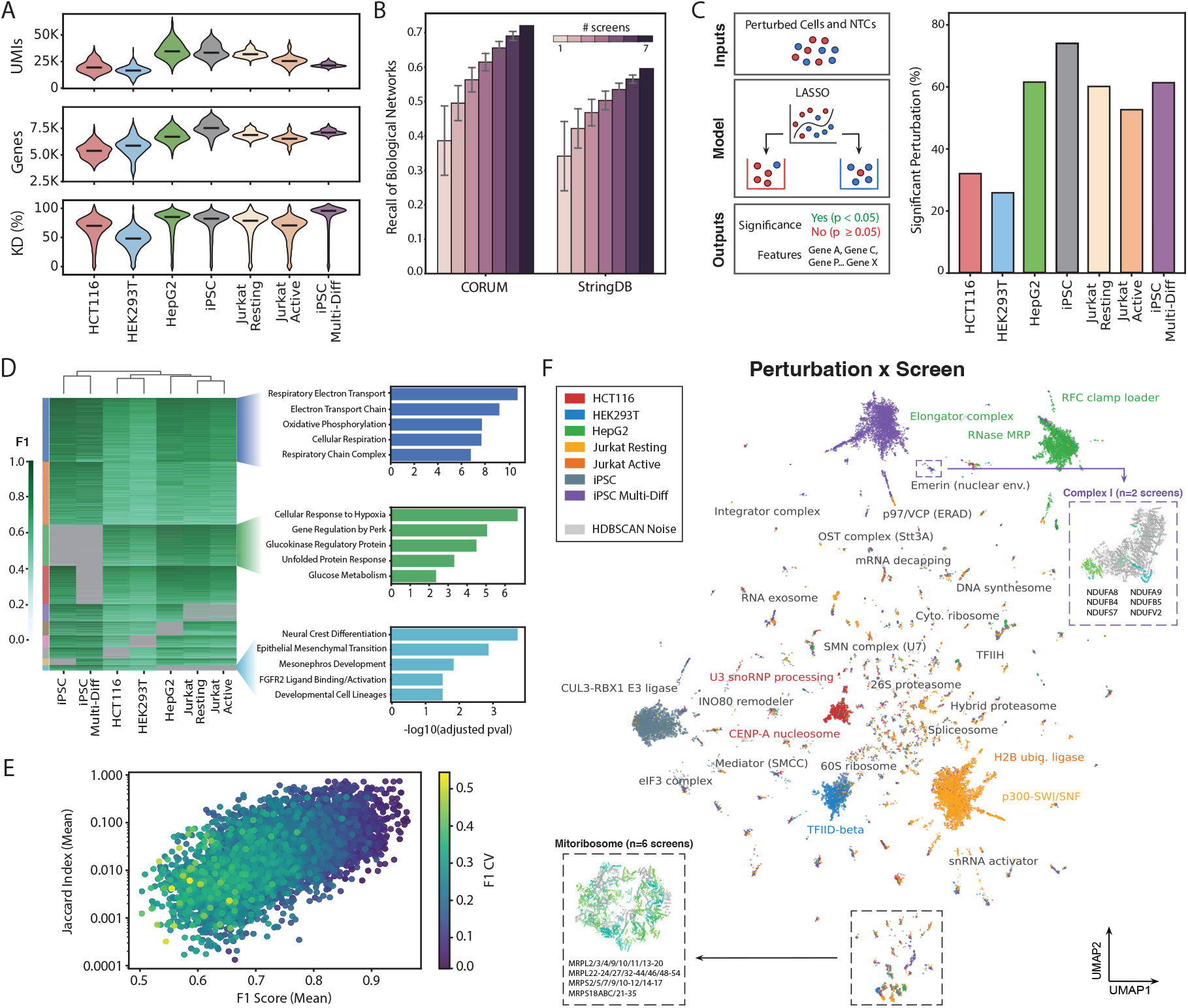
X-Atlas/Pisces enables identification of context-dependent and context-independent perturbations. **(A)** Distribution of median UMIs per cell (top), median genes per cell (middle), and median on-target KD % (bottom) across GEMs for each screen in X-Atlas/Pisces. Black lines indicate median. **(B)** Recall of annotated gene pairs from CORUM (left) and STRING (right), calculated across all possible subsets of 1–7 screens. Error bars represent standard deviation across subsets. **(C)** Schematic of binary classifier for classifying perturbed cells from non-targeting controls (NTCs) (left). Input is a balanced mixture of perturbed cells for a given perturbation and NTCs. Model is logistic regression with L1-penalization (LASSO). Outputs are significance of the perturbation as measured by a one-sided binomial test and the features (genes) used to make the prediction. Percentage of perturbed cells with a significant phenotype (FDR *<* 0.05) as measured by a LASSO binary classifier (right). **(D)** Heatmap of F1 scores for perturbations in X-Atlas/Pisces clustered based on F1 scores across screens. Rows represent perturbations (n=6,412) that pass the following filtering criteria: (1) present in ≥ 3 screens, (2) ≥ 10 cells in each screen the perturbation was present in, (3) significant phenotype (FDR *<* 0.05) in at least 1 screen, and (4) F1 score ≥ 0.75 in at least 1 screen. Grey represents absence of a perturbation in that screen. Columns represent screens that are hierarchically clustered by similarity in F1 scores (colorbar). **(E)** Mean F1 score versus mean Jaccard index for perturbations across screens. Points are colored by F1 score coefficient of variation (CV) (colorbar). **(F)** UMAP of significant perturbations with at least one non-self feature (n=35,016) colored by screen. Clusters are labeled with CORUM complex enrichments, colored black if ≤ 50% of perturbations are from any individual screen, or the majority screen color otherwise. Inset structures of representative protein complexes are color-coded by whether the protein is targeted by a genetic perturbation in the indicated clusters (green shades are present, gray is absent).

To quantify the fraction of perturbations that induced a measurable transcriptomic change, we used a LASSO binary classifier trained using an 80:20 train:test split on perturbations containing at least 10 cells with an equal number of non-targeting control cells. The fraction of perturbations with a significant transcriptomic phenotype (FDR *<* 0.05) varied substantially across screens, ranging from 25.32% in HEK293T cells to 72.80% in iPSCs (Figure 2C).

We performed hierarchical clustering of perturbations across all screens using F1 scores generated by the binary classifier. Notably, screening contexts from similar lineages clustered together, highlighting a clear structural division between conserved and context-specific effects (Figure 2D). Furthermore, individual perturbation clusters displayed specificity scores that directly reflected their degree of context dependence (Supplementary Figure 2B). For example, conserved perturbations demonstrated consistently strong effects compared to other subsets (cluster 0; blue), while perturbations highly specific to iPSCs clustered distinctly from other screens (cluster 9; teal). Pathway enrichment of perturbation subsets with similar F1 scores confirmed the observed context specificity of perturbation effects (Figure 2D). Perturbations that maintained high effect sizes across all contexts (Supplementary Figure 2B, left) were strongly enriched in shared metabolic functions, such as respiratory electron transport and oxidative phosphorylation. Conversely, perturbations with high effect sizes only in specific contexts were enriched in pathways related to canonical cell type identities. For example, a cluster with the highest F1 score in HepG2 (Supplementary Figure 2B, middle) was enriched for perturbations that modulate glucose metabolism and cellular response to hypoxia, aligning with the established profile of this hepatocellular carcinoma line as a canonical model for HIF-1-driven metabolic reprogramming in hypoxic solid tumors [44, 45]. Perturbations driving high effect sizes exclusively in iPSC contexts (Supplementary Figure 2B, right) were enriched in pathways related to cellular differentiation, including developmental cell lineages, neural crest differentiation, and epithelial-mesenchymal transition. These ontologies capture the fundamental biology of pluripotent stem cells, which undergo epithelial-mesenchymal transition during differentiation to neural crest lineages [46], further supporting the context-specificity of perturbation effects.

To assess the consistency of perturbation responses across contexts, we compared cross-screen phenotypic strength (mean F1 score) to the similarity of induced transcriptomic features (mean Jaccard index) (Figure 2E). We observed a strong positive correlation between perturbation detectability and feature conservation (Spearman=0.642, *p*-value *<* 0.001). Perturbations with lower mean effect sizes exhibited high phenotypic variability across screens and minimal feature overlap. In contrast, strongly detectable perturbations demonstrated uniform cross-screen strength and greater feature sharing. However, even among these robust, relatively context-independent perturbations, absolute feature overlap remained low, indicating that the majority of transcriptomic shifts are still driven by cell type.

To systematically map these relationships, we projected perturbations with an observable phenotype, defined as perturbations with a significant effect (*p*-value *<* 0.05) and classifier identification of at least one feature other than the direct CRISPRi target, onto a shared UMAP embedding (Figure 2F; Supplementary Figure 2C). Although context-dependent downstream effects dominate the global topology, the embedding successfully isolates tightly coherent sub-clusters characterized by varying degrees of cross-screen integration (Supplementary Figure 2D). These modules exhibit STRING interaction scores significantly higher than expected by chance (*p*-value *<* 8.44 × 10^−16^; Supplementary Figure 2E) and cluster composition reveals a spectrum of biological dependency, where perturbations that target core machinery group together across nearly all screens. For example, in the cluster associated with the mitochondrial ribosome [47], we found perturbations targeting 39S and 28S subunit components from all screens with sufficient cell numbers for feature detection. In contrast, other modules are strictly context-dependent. For example, the labeled cluster containing a subset of Respiratory Complex I components [48] is driven almost entirely by iPSC-derived perturbations. This separation may reflect intrinsic differences in metabolism across different cell types. Unlike immortalized cancer cells, which exploit Warburg-type aerobic glycolysis [49], primary cells in the pluripotent state or exiting pluripotency lack this oncogenic rewiring [50, 51] and exhibit distinctive transcriptomic responses to perturbations targeting Complex I. Taken together, these results demonstrate that we can successfully distinguish between broadly conserved biological pathways and specific regulatory programs that define unique cell states using X-Atlas/Pisces.

### 2.3 X-Cell generalizes perturbation effects across cellular contexts

X-Cell enables robust prediction of perturbation effects across diverse cellular contexts. We first evaluated the diffusion LM framework for perturbation prediction across cellular contexts with limited perturbation coverage, corresponding to a few-shot context generalization setting (Figure 3A; Section 4.1.4). Initialized from scGPT encoder weights, X-Cell was continually pre-trained on the X-Atlas/Pisces perturbation corpus spanning four cell types (HCT116, HEK293T, HepG2, and iPSC), and evaluated on an internal held-out set of 200 genes from iPSC and HepG2 in the full protein-coding gene context. The pre-trained model was subsequently fine-tuned on the Replogle–Nadig dataset [12, 52] and Parse-1M to benchmark performance on highly variable genes.

**Fig. 3.**
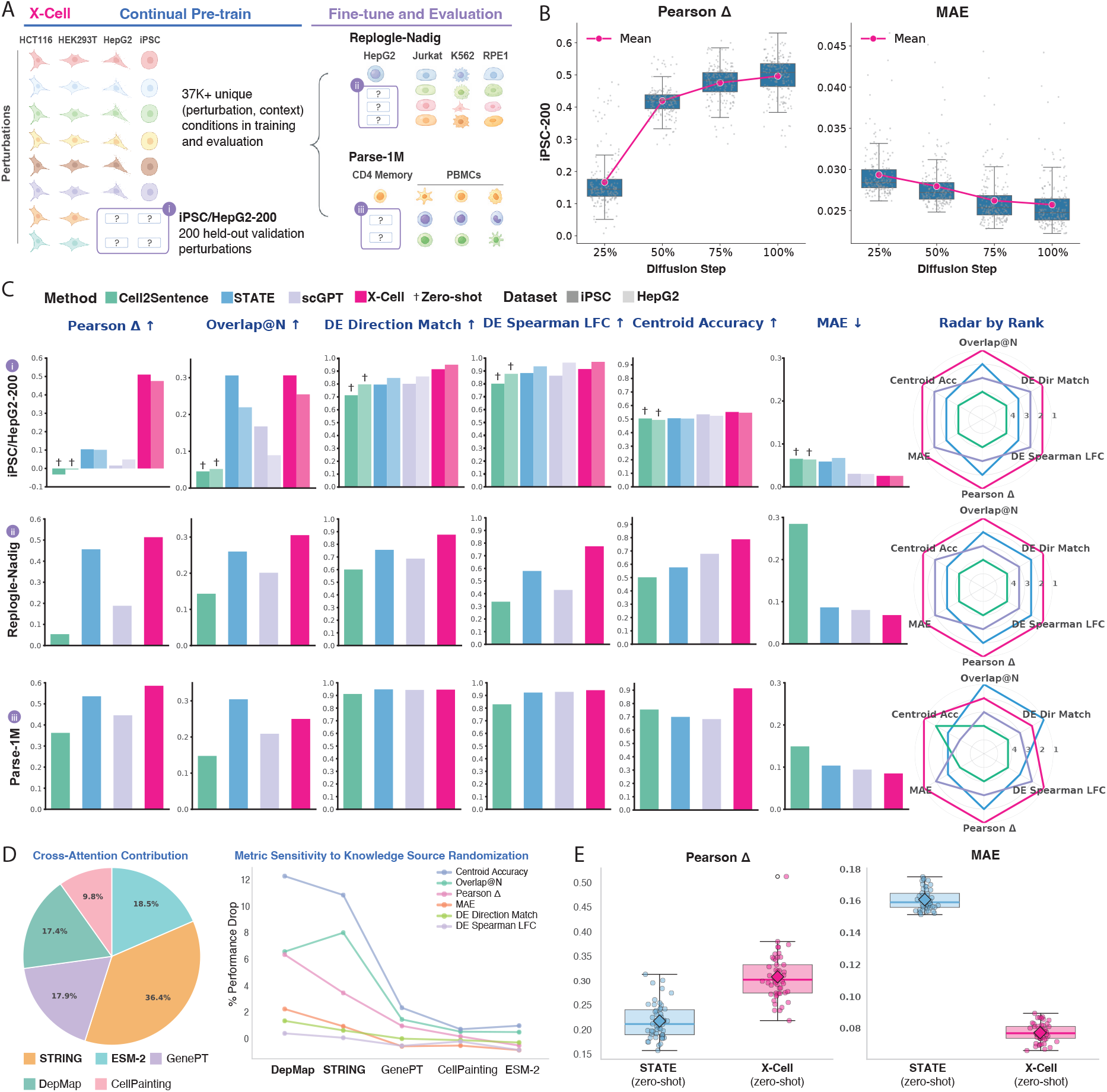
X-Cell achieves robust generalization across cellular contexts and perturbation benchmarks through diffusion pre-training. **(A)** X-Cell, initialized with scGPT encoder weights, was continually pre-trained on the X-Atlas/Pisces perturbation corpus across four screens (HCT116, HEK293T, HepG2, and iPSC), using over 10 million cells and 37K perturbation–context combinations for training and evaluation. Model performance is evaluated on the 200 held-out validation perturbations (iPSC/HepG2-200). Fine-tuning is evaluated on two external datasets: Replogle–Nadig [12, 52] and Parse-1M [53]. Replogle–Nadig includes genetic perturbations across four training cell lines and 380 HepG2 test perturbations. Parse-1M features PBMC cells with 59 ligand signaling test perturbations in CD4 memory cells. See Section 4.1.4. **(B)** Diffusion LM pre-training enables iterative refinement of predicted gene expression during inference. Line plot shows Pearson Δ and MAE across four diffusion steps on the iPSC-200 validation set. **(C)** Benchmark comparison of X-Cell with Cell2Sentence [22], STATE [27], and scGPT [8] across iPSC/HepG2-200 validation perturbations, Replogle–Nadig, and Parse-1M. Performance is evaluated using `cell-eval` metrics including Pearson Δ, Overlap@N, DE Direction Match, DE Spearman LFC, Centroid Accuracy, and MAE (Section 4.1.8). Radar plots summarize the relative ranking of the four benchmark models. **(D)** Prior knowledge attribution on Replogle–Nadig test perturbations. Pie chart shows average attention weights from X-Cell’s final cross-attention layer. Line plot shows the performance drop on six metrics after randomizing embeddings from each knowledge source. **(E)** Zero-shot benchmark of X-Cell on a Tahoe-100M subset containing single-target inhibitor drugs at mid dosage, compared with STATE (Section 4.1.7). Each point represents the aggregated metric across 12 drugs in each of 50 cell lines. Box plots show the median and interquartile range across cell lines; diamonds denote the mean.

Diffusion LM pre-training enables iterative refinement of predictions at inference time by remasking a proportion of predicted gene expression values and reconditioning on the control transcriptome at each generation step. On the held-out iPSC-200 perturbations, X-Cell predictions progressively aligned with observed transcriptional responses, with increasing Pearson correlation of log-fold change (Pearson Δ) and decreasing mean absolute error (MAE) across diffusion steps (Figure 3B; Section 4.1.8). These findings highlight diffusion-driven in-context refinement as a key mechanism enabling X-Cell to adapt and improve perturbational predictions during inference.

We benchmarked the performance of X-Cell against several state-of-the-art perturbation prediction models (Cell2Sentence (C2S) [22], STATE [27], and scGPT [8]) for the iPSC and HepG2-200 validation sets across multiple `cell-eval` metrics [27]: fold-change concordance (Pearson Δ), differential expression accuracy (Overlap@N, DE Direction Match, DE Spearman LFC), perturbation specificity (Centroid Accuracy), and overall predictive fidelity (MAE) (Section 4.1.8). Due to compute constraints to retrain all benchmark models at X-Atlas/Pisces scale, we report zero-shot performance of the C2S 2B model, and a STATE (SE+ST) model trained on a substantial subset (5.5M cells) of X-Atlas/Pisces data from the matching four cell lines (Section 4.1.4). X-Cell outperforms all benchmark methods (Figure 3C, row i), achieving Pearson Δ values of 0.51 and 0.48, both more than five-fold greater than the next best model, STATE (0.10), highlighting the impact of architectural advances beyond data scale (Supplementary Table B7).

We then evaluated X-Cell’s generalizability to external CRISPR perturbation datasets by fine-tuning the model on the Replogle–Nadig dataset [12, 52] (Section 4.1.7), which introduces a distributional shift towards essential gene knockdowns across four cell lines (K562, RPE1, HepG2, and Jurkat) and 2,000 highly variable genes. Comparing model performance on 380 held-out genes from the HepG2 cell line, X-Cell ranks first for all six reported metrics and achieves the highest predictive specificity, with a Centroid Accuracy of 0.79 (Figure 3C, row ii; Supplementary Table B7). All benchmark models were fine-tuned and evaluated using the same data split (Section 4.1.4). To investigate which biological priors drive these predictions, we first examined cross-attention weights across knowledge sources, revealing STRING and ESM-2 as the dominant knowledge sources influencing model predictions (Figure 3D, Section 4.1.10). Because X-Cell is initialized with scGPT gene embeddings, which are further updated during training, we exclude scGPT from this attribution analysis. Further gene set enrichment analysis (GSEA) on gene rankings derived from attention weights for each prior knowledge source showed that STRING-based rankings produced the strongest enrichment in MSigDB Hallmark pathways (Supplementary Figure 4). We then quantified task-specific contributions using a knowledge source randomization analysis, in which pre-trained embeddings from each of the five prior knowledge sources were independently randomized to report the resulting percentage drop in performance. DepMap and STRING led to the largest performance degradations, particularly in specificity and DEG prediction accuracy, indicating that these priors provide the strongest task-relevant signals for differentiating the effects of perturbations (Figure 3D).

We further evaluated X-Cell’s ability to transfer knowledge from genetic perturbations to signaling and chemical perturbations. In a fine-tuning setting, we incorporated ligand information via cross-attention using the Parse-1M dataset [53], achieving strong performance on a held-out set of 59 signaling perturbations (Figure 3C, row iii; Section 4.1.7). Next, we tested X-Cell’s generalization to predicting the effects of drug perturbations in a zero-shot setting. We constructed a subset of the Tahoe-100M dataset containing inhibitor drugs with single-target mechanisms of action (MOA), establishing a direct link between genetic perturbations and drug effects (Section 4.1.7). In this setup, X-Cell encodes the MOA gene as the perturbed target and the control cell-line transcriptome as input to predict drug-induced expression responses (Figure 3E). On zero-shot tasks, we benchmarked against a STATE model trained on 3.1M cells from public datasets only (Section 4.1.5) [12, 52, 55]. Despite never observing Tahoe perturbations during training, X-Cell achieves stronger performance in both Pearson Δ (mean 0.31 versus 0.22) and MAE (mean 0.08 versus 0.16). A stratified analysis further suggests that predictions are more effective for cell lines that are similar to the X-Cell training corpus, highlighting the importance of biologically relevant training contexts for successful cross-modality generalization (Supplementary Figure 6). These results highlight X-Cell’s ability to generalize across perturbation modalities and suggest a path toward integrating perturbation modeling with drug discovery.

### 2.4 X-Cell predicts T cell inactivating perturbations in a zero-shot manner

To nominate context-dependent perturbations, we analyzed genome-scale Perturb-seq data in Jurkat cells across resting and active states. We focused on perturbations critical for T cell activation, which we hypothesized would exhibit state-dependent effects. To investigate the transcriptional shifts associated with T cell state transitions, we defined an “inactivation index” to quantify the degree to which a specific perturbation shifts an activated cell state toward a resting phenotype; a higher index indicates a global transcriptome profile that more closely resembles unperturbed resting cells (Figure 4A; Section 4.1.9). As expected, members of the *CD3* complex (*CD3D, CD3E, CD3G*, and *CD247*) and their downstream effectors (*ZAP70, LCP2, CDK6*, and *ITK*) exhibited high inactivation indices. We also identified several putative T cell inactivators—including *IRAK4, CHD5, LRBA, LIMS2, BRCA1, APPL2*, and *WDR53* —noting that some may represent Jurkat-specific dependencies.

**Fig. 4.**
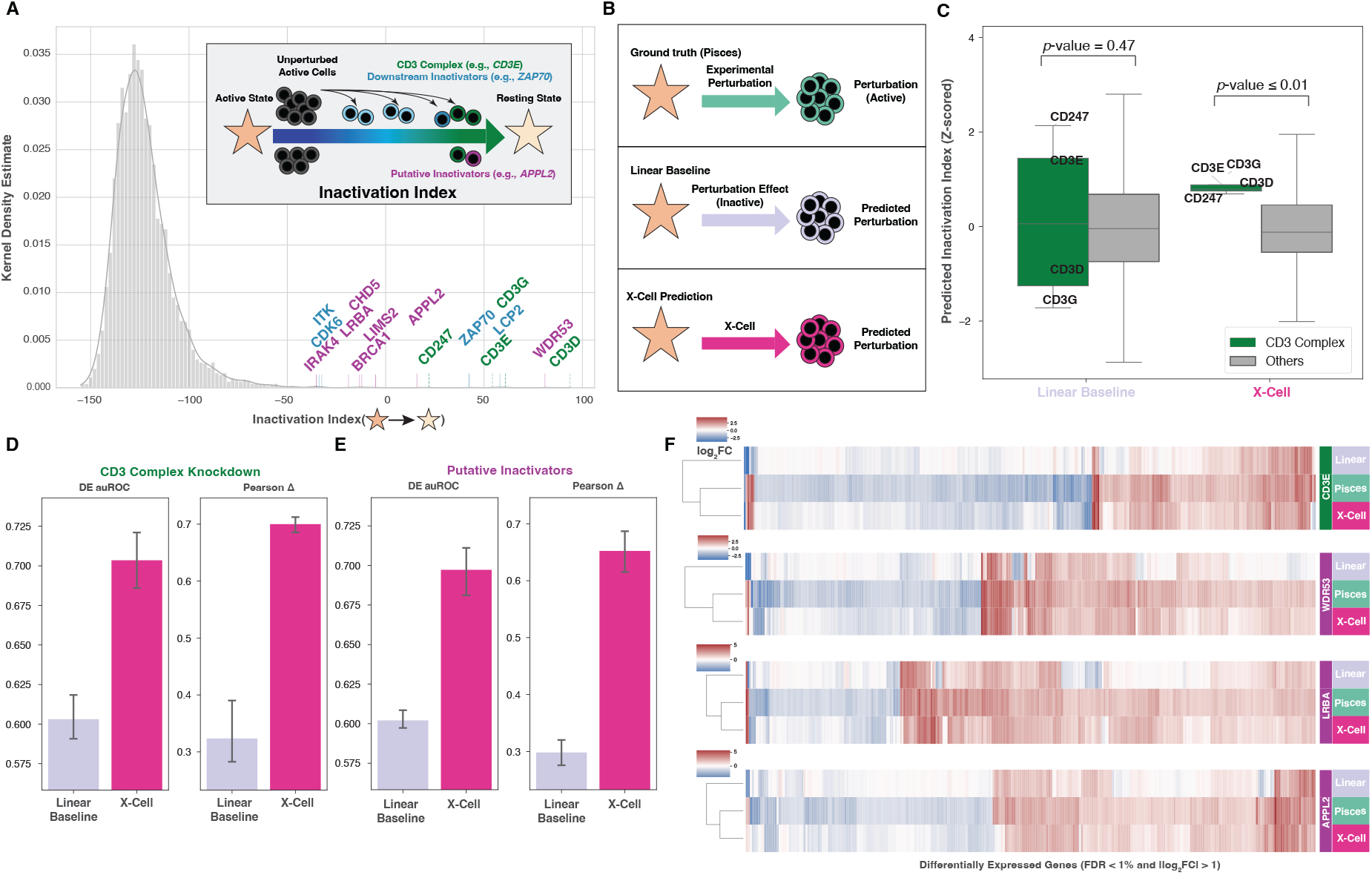
X-Cell distinguishes physiological state transitions in Jurkat T cells. **(A)** Schematic representation of the inactivation index. Stars represent non-targeting controls (active or resting), while circles represent perturbations. The inactivation index quantifies the shift of activated cells toward a resting phenotype based on global transcriptome similarity. **(B)** Schematic of the ground-truth (Pisces atlas dataset), linear baseline, and X-Cell predictions. In the linear baseline, the transcriptional impact (Δ) of a perturbation measured in resting Jurkat cells is additively applied to the log-normalized profile of unperturbed active Jurkat cells. **(C)** Boxplots show the distribution of inactivation indices (Z-scores) for validated T cell activation regulators, the CD3 complex (green, including *CD3D, CD3E, CD3G*, and *CD247*), compared to 4,455 other perturbations (gray). X-Cell predicts a significant shift toward the resting state compared to the linear baseline (Mann-Whitney U, *p*-value *<* 0.01). **(D)** Performance comparison between X-Cell and the linear baseline for the CD3 complex. Evaluation metrics include auROC for identifying differentially expressed genes (DEGs) and Pearson Δ metrics for predicting the magnitude of transcriptional changes. Barplot shows average score and error bars indicate 95% confidence intervals. **(E)** Similar to (D), showing X-Cell’s capability in identifying transcriptional changes of putative inactivators (*WDR53, APPL2*, etc.). **(F)** The heatmaps show log_2_ fold change in expression values and clustering of predicted transcriptional profiles for *CD3E, WDR53, LRBA*, and *APPL2*, demonstrating that X-Cell predictions closely align with ground-truth Pisces atlas data.

After establishing these biological benchmarks, we evaluated X-Cell’s ability to predict those perturbations’ effects in a zero-shot manner. We fine-tuned X-Cell on resting Jurkat datasets from X-Atlas/Pisces, and compared its performance against a linear baseline, which transfers the perturbation impact from the Jurkat resting datasets to Jurkat active control cells (Figure 4B). Compared to a linear baseline, X-Cell predicted a significantly more pronounced and accurate shift toward the resting state for members of the *CD3* complex (Mann-Whitney U, *p*-value *<* 0.01; Figure 4C). Furthermore, X-Cell outperformed the linear baseline in predicting both the identity of differentially expressed genes (DEGs), as measured by auROC (Section 4.1.8), and the magnitude of transcriptional changes, as measured by Pearson Δ metrics, for both the *CD3* complex (Figure 4D) and putative inactivators (Figure 4E). Hierarchical clustering of predicted transcriptional profiles for *CD3E, WDR53, LRBA*, and *APPL2*, showed that X-Cell’s predictions align significantly more closely with ground-truth X-Atlas/Pisces data than the linear baseline, demonstrating the model’s superior ability to generalize out-of-context (Figure 4F).

To elucidate the mechanistic basis of this performance, we analyzed the model’s cross-attention scores, hypothesizing that external knowledge bases provide critical, context-specific biological priors during inference. Specifically, during the simulation of *CD3E* perturbation in active Jurkat cells, we observed a marked increase in the attention weights assigned to ESM-2, GenePT, and DepMap embeddings. This focus is biologically corroborated by the significant enrichment of the top 1,000 genes prioritized by these modalities in the KEGG TCR signaling pathway (hsa04660) [56]. These results suggest that X-Cell dynamically leverages relevant external knowledge to refine its predictions within specific biological contexts (Supplementary Figure 5).

### 2.5 X-Cell-Ultra follows universal neural scaling laws, charting a path to further performance gains by increasing compute and data

Encouraged by X-Cell’s ability to generalize to unseen exposures, we next investigated the effects of scaling along both the data and model parameter axes. On the data axis, we leveraged the majority of X-Atlas/Pisces, featuring *>*20 million cells and *>*152,000 unique perturbation–context combinations for training and evaluation, across GEM-X and FLEX platforms (Figure 5A). On the modeling axis, building on anti-collapse architectural innovations validated in our ablation studies (Supplementary Figure 7A; Supplementary Table A1), we scaled X-Cell to 4.9 billion parameters, *X-Cell-Ultra*, and conducted a systematic scaling analysis to determine whether perturbation prediction benefits from the same scaling principles that have driven progress in large language models. We then deployed X-Cell-Ultra for genome-scale perturbation inference in unseen biological contexts (Figure 5A). As the biological foundation model with the largest parameter count to date, X-Cell-Ultra achieves ∼ 41% model FLOPs utilization (MFU) on 64–128 H200 GPUs, comparable to production-grade LLMs (Supplementary Figure 7B; Section 4.1.3). Notably, without relying on scGPT pre-trained weights, X-Cell-Ultra departs from the conventional paradigm of pre-training on observational data and fine-tuning on perturbations, instead representing the first perturbation foundation model trained entirely on large-scale interventional data (Figure 5A).

**Fig. 5.**
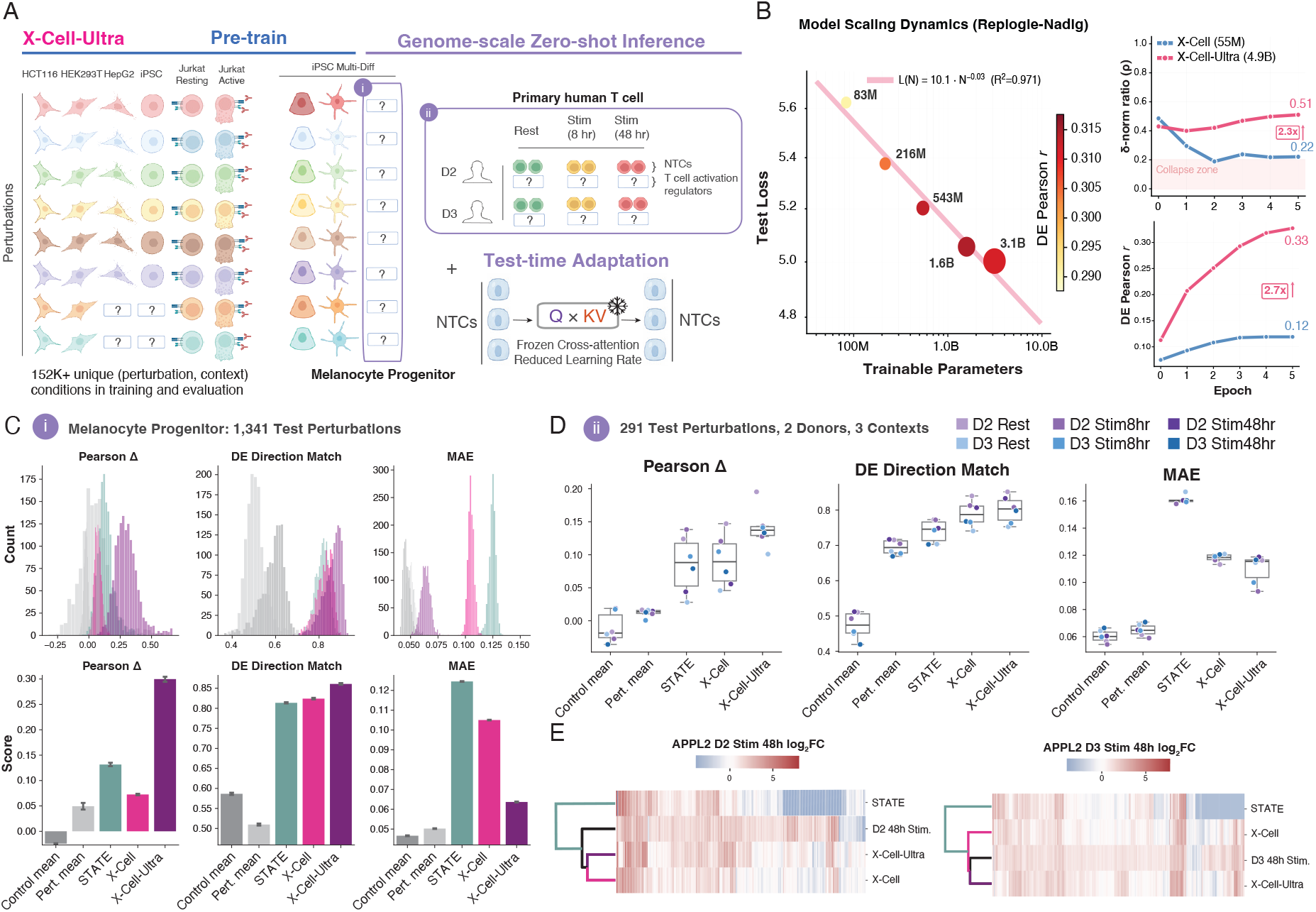
X-Cell-Ultra generalizes to melanocyte progenitors and primary human T cells in a zero-shot manner. **(A)** Schematic shows pre-training of X-Cell-Ultra on X-Atlas/Pisces datasets, excluding the melanocyte progenitor population from the iPSC Multi-Diff screen. We evaluated X-Cell-Ultra in a zero-shot manner on the melanocyte progenitor population as well as primary human T cell datasets from Zhu et al. [57]. We used test-time adaptation to calibrate the models on these datasets using self-supervised objectives. **(B)** Test loss follows a power law with trainable parameters (*L*(*N*) ∝ *N* ^−0.03^, *R*^2^ = 0.97; Table A4). Five models (83M–3.1B) were trained for 20 epochs on the Replogle-Nadig dataset under identical hyperparameters. Circle size is proportional to parameter count; color indicates DE Pearson *r*. Line plots (right) show how X-Cell-Architecture (blue) and X-Cell-Ultra-Architecture (pink) compare with respect to collapse metrics *δ*-norm ratio (top) and Pearson correlation of differentially expressed genes (bottom) on test set. **(C)** Performance of the control mean baseline, perturbation mean baseline, STATE, X-Cell, and X-Cell-Ultra with respect to Pearson Δ (left), differential expression direction match (middle), and mean absolute error (right). Histogram (top) shows the metrics within 1,341 perturbations and the barplots (bottom) show mean ± 95% C.I. **(D)** Boxplots show the distribution of Pearson Δ (left), differential expression direction match (middle), and mean absolute error (right) for 291 perturbations in resting and stimulated (2 time points) primary human T cells for 2 different donors. **(E)** Heatmaps show the log_2_ fold change in expression of APPL2 in D2 (left) and D3 (right) 48 hours post-stimulation as observed by the data and predicted by STATE, X-Cell, and X-Cell-Ultra.

To rigorously evaluate whether the scaling and training advances embodied by X-Cell-Ultra translate into genuine biological generalization, we assessed zero-shot performance on two cell populations entirely absent from training: melanocyte progenitors from our iPSC Multi-Diff screen and primary human CD4^+^ T cells from an independent study [57] (Figure 5A; Section 4.1.4). In both cases, we applied test-time adaptation (TTA; Section 4.1.3) (self-supervised fine-tuning on unlabeled non-targeting control cells from the target domain using only the MMD loss) to calibrate the model’s self-attention representations to the target expression manifold before perturbation inference.

We first performed genome-scale inference on held-out melanocytes from the iPSC Multi-Diff screen, predicting responses to 1,341 genetic perturbations (Section 4.1.7). X-Cell-Ultra outperforms the control mean baseline, perturbation mean baseline, STATE, and X-Cell across Pearson Δ, DE Direction Match, and MAE (Figure 5C). The gain over X-Cell is particularly pronounced for Pearson Δ and DE Direction Match, indicating that the additional model capacity specifically improves the recovery of perturbation-specific transcriptional signatures rather than merely reducing reconstruction error.

To test generalization beyond cell lines to primary human cells (Section 4.1.7), we evaluated on 291 perturbations of T cell regulators from primary CD4^+^ T cells across two donors in resting and stimulated states (8 and 48 hours post CD3/CD28 stimulation) [57]. These cellular contexts are completely absent from the X-Atlas/Pisces training corpus and represent donor-specific and time-dependent activation programs. X-Cell-Ultra achieves higher Pearson Δ and DE Direction Match than both X-Cell and STATE across all donor–condition combinations, with particularly pronounced gains in the stimulated conditions where context-dependent regulatory rewiring is most extreme (Figure 5D). As a case study, *APPL2* –a putative T cell inactivator identified in our Jurkat analysis (Section 2.4) shows that X-Cell-Ultra accurately recapitulates the stimulated-state transcriptional signatures among both donor samples (Figure 5E), providing independent confirmation of the model’s biological fidelity across experimental systems.

To characterize scaling behavior, we trained five model variants spanning 83M to 3.1B parameters for 20 epochs each under identical hyperparameters (Supplementary Table A4; Supplementary Figure 7C– E). Training loss follows a power law *L*(*N*) ∝ *N*^−*α*^ with *α*_train_ = 0.32 (*R*^2^ = 0.96), squarely within the range reported for large language models (*α* ≈ 0.076–0.34) [58, 59]. Test loss decreases monotonically from 5.62 (83M) to 5.00 (3.1B), with a shallower exponent (*α*_val_ = 0.03, *R*^2^ = 0.97) (Figure 5B). Downstream biological metrics also improve with scale: DE Pearson *r* increases from 0.291 (83M) to 0.315 (1.6B), an 8% gain in fold-change correlation, with the 3.1B model sustaining *r* = 0.312. Per-perturbation MAE decreases monotonically across all model sizes (Supplementary Figure 7D).

To interpret the divergence between train and test scaling exponents, we note that all five models were trained on identical data (20 epochs, ∼ 5,300 unique perturbation–context groups). The Chinchilla decomposition [59],

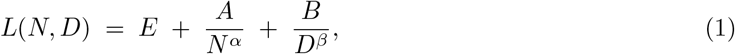

separates loss into irreducible noise (*E*), model-capacity (*A/N* ^*α*^), and data-starvation (*B/D*^*β*^) terms. Because *D* is held constant across our scaling sweep, the data-starvation contribution *E* + *B/D*^*β*^ acts as a floor independent of model size. The steep train exponent (*α*_train_ = 0.32, within the range reported for large language models [58, 59]) shows that the capacity term *A/N*^*α*^ drives training loss reduction and demonstrates that larger models with enhanced capacity learn perturbation prediction more efficiently. The shallow test exponent (*α*_val_ = 0.03) shows that the data-starvation floor dominates generalization such that adding parameters beyond ∼ 1.6B does not reduce the test loss appreciably, and DE Pearson *r* plateaus at *r* ≈ 0.31 (Supplementary Figure 7E). Because the natural data unit for set-level MMD models is a unique (perturbation, context) group rather than an individual token, the precise compute-optimal allocation *D*^∗^(*N*) for this modality remains an open question that will require controlled iso-FLOP experiments varying both *N* and *D* independently. Nonetheless, the expansion in unique conditions from Replogle-Nadig (∼ 5,300) to the full X-Atlas/Pisces compendium (∼ 151,000) directly motivated the deployment of the larger 4.9B X-Cell-Ultra. These results establish, to our knowledge for the first time, that single-cell perturbation prediction exhibits power-law scaling consistent with frontier language models, while revealing that the diversity of unique perturbation–context combinations, rather than token count or model capacity, is a current binding constraint on downstream generalization performance.

## 3 Discussion

A primary motivation for modeling causal interventional dynamics is their potential to accelerate therapeutic discovery, from target identification and validation to patient-specific prioritization [37]. To tackle the fundamental challenge of predicting context-dependent gene regulatory responses, we developed X-Atlas/Pisces — the largest genome-wide CRISPRi Perturb-seq resource to date — and the X-Cell diffusion language model. By coupling its training on the diverse, multi-context Pisces dataset with the active integration of prior knowledge, including coherent literature embeddings alongside biochemical, essentiality, and morphological data, X-Cell infers how regulatory networks rewire across contexts. This powerful combination enables the model to predict perturbation-specific differentially expressed genes with high fidelity, unlocking advanced counterfactual inference in diseased human cells. Ultimately, X-Cell’s ability to capture divergent transcriptional responses across distinct states presages a future where the applicability of therapeutic hypotheses can be simulated across the entire diversity of human cellular contexts.

One striking demonstration of this context-aware predictive power is the zero-shot prediction of T cell inactivation in stimulated Jurkat cells. By fine-tuning only on perturbations from the resting state, X-Cell successfully imputed the distinct, state-specific transcriptional consequences of perturbing the canonical CD3 complex upon T cell receptor activation with CD3/CD28 antibodies. Further, the model generated high-fidelity predictions for putative regulators, identifying *APPL2* and *LRBA* as potent T cell inactivators. Importantly, these zero-shot predictions are not only supported by established mechanistic literature in distinct cellular contexts, but were directly validated by external ground-truth data. LRBA regulates lysosomal degradation of a co-inhibitory receptor CTLA4 [60], with its depletion accelerating CTLA4 turnover. APPL2 is a Rab5-associated endosomal adaptor that drives F-actin remodeling [61], suggesting it may link endosomal trafficking to the cytoskeletal dynamics required for TCR signaling, functionally similar to RASA2 [62]. During the preparation of this manuscript, *APPL2* was independently recovered as a putative regulator in a genome-wide Perturb-seq screen in primary human CD4^+^ T cells, while *LRBA* was identified as a downstream target of a stimulation-dependent STAT3 regulatory network [57]. Resolving these underlying mechanisms will require further experimental validation.

The robustness of X-Cell’s performance extends beyond leveraging prior knowledge to predict the effects of perturbations in cell states unseen in cell types that were seen in training. Crucially, X-Cell-Ultra demonstrates zero-shot generalization to entirely unseen cell lineages — a critical threshold that previous foundation models have yet to cross. When evaluated on melanocyte progenitors from our iPSC Multi-Diff screen (a cell type completely absent from the training corpus), X-Cell-Ultra yielded highly accurate perturbation predictions, excelling specifically in the recovery of differentially expressed genes. This indicates that the model is not merely interpolating within the distribution of its training data, but fundamentally learning how the manifold of gene expression transforms in novel biological contexts. X-Cell also bridges the translational gap between tractable laboratory cell lines and complex primary human tissues. When evaluated on primary human CD4^+^ T cells from multiple donors, X-Cell-Ultra successfully predicted highly accurate, context-specific perturbation responses. We note that this strong performance is driven in part by the inclusion of genome-scale Jurkat perturbation data in our training corpus, as this is a classic and well-validated model system for studying the T cell receptor signaling pathway [63]. This successful translation highlights a powerful paradigm for applying perturbation foundation models to drug discovery: generating pre-training data in tractable experimental systems with high relevance to therapeutically critical human cell types, ultimately allowing us to computationally “screen the unscreenable” in primary patient cells.

Fundamentally, our analyses reveal that perturbation prediction performance demonstrably improves as model capacity scales. By training X-Cell up to 4.9 billion parameters, we observed measurable gains in predictive fidelity, validating that biological foundation models benefit from the scaling principles established in natural language processing. We hypothesize that the synergistic combination of further architectural refinements and a massive expansion of contextually diverse, large scale perturbation datasets will serve as a catalyst for improving performance of perturbation foundation models. Pushing these boundaries is likely the prerequisite for unlocking emergent properties capable of simulating complex biological phenomena in unseen cellular contexts to a level that enables high accuracy identification of novel therapeutic hypotheses.

Even with increased scale, generalizing globally across all diverse cell types remains a profound challenge. As anticipated, and as directly evidenced by our perturbation embedding, the transcriptomic consequences of identical genetic interventions can be heavily influenced by biological context. We speculate that the core, shared features that drive the cross-screen clustering of members of macromolecular complexes may represent the ubiquitous, proximal consequences of a genetic perturbation, whereas the cell-type-specific features reflect cascading, context-specific downstream consequences. Because of this context-specificity, simple heuristic baselines — such as predicting the mean perturbation effect within a given context — are stubbornly difficult to outperform consistently across all evaluation metrics. Indeed, claiming absolute superiority over baselines often masks intrinsic trade-offs; models can excel at minimizing absolute error by being overly conservative but miss key changes in gene expression that represent the hallmark effects of a perturbation in a novel context.

Importantly, while other approaches struggle to accurately predict specific differentially expressed genes (DEGs) over simple mean shifts, X-Cell excels in this domain. X-Cell does so not by optimizing a single metric at the expense of others, but by mitigating conservative collapse, a common failure mode where models correctly predict the direction of differential expression but severely suppress its magnitude. The combination of our contextually diverse X-Atlas compendium and rich prior knowledge embeddings provides the necessary anchor for the model to begin generalizing beyond these simpler baselines, accurately predicting the direct and indirect biological sequelae of experimental interventions. While the precise internal mechanics of deep learning models remain inherently opaque, our ablation studies strongly suggest that X-Cell bridges these divergent physiological states by dynamically attending to multi-modal prior knowledge embeddings.

Our scaling analysis reveals that while training loss follows the same power-law dynamics established by the Chinchilla framework for large language models, downstream biological metrics saturate before the largest models are fully utilized (Section 2.5). On the Replogle-Nadig dataset, fold-change correlation plateaus above ∼1.6B parameters — a consequence of the ∼5,300 unique (perturbation, context) groups bounding effective diversity below what additional parameters can exploit. This data-limited regime motivated our deployment of the 4.9B X-Cell-Ultra on the substantially larger X-Atlas/Pisces compendium, where more unique conditions provide greater biological diversity. More broadly, these results indicate that expanding contextually diverse perturbation compendia in proportion to model scale is the highest-leverage path to future gains, and that resolving the precise compute-optimal frontier for set-level biological models will require controlled iso-FLOP experiments that independently vary model size and dataset diversity.

Despite achieving state-of-the-art performance, X-Cell also highlights critical frontiers that remain in perturbation modeling. Our high fidelity perturbation predictions are currently constrained to isolated, single-cellular contexts. For therapeutic discovery, the true consequence of a perturbation is rarely confined to a single cell type; it cascades through tissue ecosystems via cell-cell interactions. While X-Cell provides immediate utility for conducting *in silico* screens to prioritize targets in cell types with known roles in disease biology, realizing the full potential of drug discovery requires predicting downstream consequences across a broader cellular context [64]. Future foundation models will need to bridge the architectural gap between precise single-cell predictions and multi-cellular tissue dynamics, necessitating training corpora that integrate spatial and systems-level perturbation data from disease-relevant models.

Collectively, this work represents a meaningful step forward in the pursuit of a generalizable perturbation foundation model. By demonstrating that discrete diffusion architectures can successfully integrate biological priors to predict complex, zero-shot physiological transitions, X-Cell underscores the immense potential of pairing generative AI with large-scale, causal data generation. As experimental modalities continue to expand to capture multi-cellular and whole tissue contexts, biologically aware modeling frameworks such as X-Cell will be well-positioned to evolve alongside them, advancing the field toward accurate *in silico* experimentation.

## 4 Methods

### 4.1 X-Cell Methodology

#### 4.1.1 X-Atlas/Pisces Training Corpus

For model training, we applied additional data filters to X-Atlas/Pisces to construct the training corpus: (1) keeping cells with the expected dual-guide pair and passing UMI and gene count filters within each screen (UMI and gene count thresholds were manually determined for each GEM), (2) filtering for genes targeting expressed genes in both HCT116 and HEK293T screens (for definition of expressed genes see Section 4.2.11), (3) exclusion of 200 validation perturbations from the iPSC and HepG2 screens (Section 4.1.7), and (4) exclusion of perturbations targeting melanocyte progenitors, which were held out for zero-shot evaluation.

Following these filters, the X-Cell training corpus consisted of 10,165,929 cells and 36,816 unique perturbation–context conditions from four screens (HCT116, HEK293T, HepG2, and iPSC). The full X-Cell-Ultra corpus contained 20,281,764 cells and 151,180 unique perturbation–context conditions from seven screens (HCT116, HEK293T, HepG2, iPSC, Jurkat Resting, Jurkat Active, and iPSC Multi-Diff). For X-Cell-Ultra training, in phase 1, high-effect initialization, we selected the top 5% highest-effect perturbations within each cell type. Candidate perturbations were first restricted to those with detectable transcriptomic effects as identified by binary classifier (FDR *<* 0.05), and subsequently ranked by the number of significant features identified from LASSO regression (Section 4.2.13). This high-effect subset consisted of 3,797,171 cells and 4,996 unique perturbation–context conditions across all screens for phase 1 training. In phase 2, training proceeded on the full corpus.

During training, target perturbations were further restricted to protein-coding genes to align with ESM2 prior knowledge keys (Section 4.1.2). After this step, the final training sets contained *>*10 million cells for X-Cell and *>*20 million cells for X-Cell-Ultra. Gene expression values were stored as raw counts and normalized during training using CP10K log1p normalization (Section A.10.2).

#### 4.1.2 Prior Knowledge Sources

To incorporate prior biological knowledge, we integrated embeddings capturing complementary representations of gene function across multiple biological levels, including language, structural, and functional information about the perturbed gene. Through cross-attention (Section 4.1.3), these priors dynamically condition perturbation predictions on a unified representation of gene function across biological modalities:

##### GenePT (text embeddings)

GenePT embeddings encode semantic representations of gene function by applying large language model text embeddings to NCBI gene summaries and related annotations. These embeddings capture functional relationships between genes learned from biomedical literature and textual gene descriptions [19]. We used the GenePT embedding release ^1^ generated using the `text-embedding-3-large` model, specifically `GenePT_gene_protein_embedding_model_3_text.pickle`.

##### ESM-2 (protein language model)

We incorporated protein sequence embeddings from the ESM-2 protein language model [38] for protein-coding genes, which captures structural and biochemical properties of proteins directly from amino acid sequences.

##### STRING (interaction network)

Protein interaction priors were obtained from embeddings derived from the STRING protein–protein interaction (PPI) network [41]. We used the STRING network embeddings ^2^ released by the SPACE project [40], where interaction networks are converted into continuous node embeddings using graph embedding approaches.

##### DepMap (gene dependency)

Functional genomics priors were derived from the DepMap CRISPR gene dependency dataset [42], which systematically measures the fitness impact of gene knockouts across hundreds of human cancer cell lines. We used the gene effect matrix `CRISPRGeneEffect.csv` from the DepMap 24Q4 public release^3^, which provides Chronos-inferred gene effect scores from pooled CRISPR screens. These embeddings capture how perturbations of specific genes influence cellular growth and survival.

##### JUMP Cell Painting (morphological phenotype)

We incorporated morphological embeddings derived from the JUMP Cell Painting dataset [43], which summarizes image-derived cellular morphology features measured in large-scale CRISPR perturbation experiments. Specifically, we used the batch-corrected embeddings from a CRISPRi screen performed in the U2OS bone cancer cell line [65]. These embeddings are derived from over 3,000 extracted morphological features, with Harmony batch correction and PCA dimensionality reduction. The resulting embeddings provide a phenotypic representation of cellular responses to gene perturbations based on high-content imaging. The processed embeddings were retrieved from the Cell Painting Gallery release^4^ using the batch-corrected feature matrix `profiles_wellpos_cc_var_mad_outlier_featselect_sphering_harmony_PCA_corrected.parquet`.

##### scGPT (transcriptomic foundation model)

We incorporated gene embeddings from the `scGPT_human` model [8] through encoder weight initialization in X-Cell, which encode transcriptomic relationships learned from large single-cell expression atlases. Because the `nn.Embedding` layer weights are updated during model training, this prior remains soft and adaptive rather than fixed like the other knowledge sources.

#### 4.1.3 Model Architecture

X-Cell is a set-level diffusion transformer that predicts perturbed gene expression profiles from control cell populations. Unlike autoregressive single-cell foundation models [8, 22], X-Cell operates on *sets* of cells and is trained explicitly on interventional data via distribution-matching objectives.

##### Input Representation

Each training sample consists of a set of *S* = 64 cells, each profiled across *G* genes (*G* ≤ 19,408 protein-coding genes after subsampling to context length *G*^*′*^ ≤ 4,000) per minibatch. For each gene position *g*, the model receives a learned gene identity embedding **e**_*g*_ ∈ ℝ^*d*^ from a shared vocabulary (followed by LayerNorm), together with a continuous value encoding *ϕ*(*x*_*g*_) ∈ ℝ^*d*^ in log1p CP10K obtained by projecting the scalar expression value through a two-layer MLP with ReLU activation and LayerNorm.

For diffusion-style training, X-Cell employs a discrete masking process inspired by masked diffusion language models [66, 67]. Let **x**^ctrl^, **x**^pert^ ∈ ℝ^*G*^*′* denote the control and perturbed expression profiles. For each sample, a replacement fraction

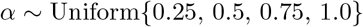

is sampled, and a subset of gene indices ℐ ⊂ {1, …, *G*^*′*^} with |ℐ| = ⌊*αG*^*′*^⌋ is selected uniformly without replacement. This defines a binary perturbation mask **M**_pert_ ∈ {0, 1}^*G*^*′* :

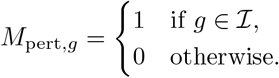

In training, the observed input expression profile is constructed by replacing control values with *groundtruth* perturbed values at the selected positions,

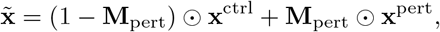

where ⊙ denotes elementwise multiplication. The binary mask is separately embedded through a learned mask embedding table *ψ*(*M*_pert,*g*_) ∈ ℝ^*d*^, which indicates whether each gene position contains control or revealed perturbed signal.

The final input token representation for gene position *g* is then given by

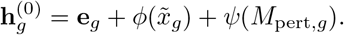

A learned `[CLS]` token **c** ∈ ℝ^*d*^, independent of the gene, value, and mask encoders and initialized from 𝒩 (0, 0.02), is prepended at position 0 to aggregate global cell-level information. The model is trained to reconstruct the full perturbed expression profile **x**^pert^ from the partially revealed input across varying replacement levels.

##### Transformer Backbone

X-Cell has 12 layers and an embedding space dimension of 512 (*L* = 12, *d* = 512, 55M parameters) and initializes gene embeddings and transformer weights from pre-trained scGPT [8], using the original Post-LN architecture with standard ReLU FFN (feed-forward network) and 4 cross-attention layers. For scalability to billions of parameters, X-Cell-Ultra (*L* = 34, *d* = 2560, 4.87B parameters) adopts a modernized architecture following LLaMA [68]: Pre-LN [69] with RMSNorm [70], SwiGLU feed-forward networks [71] with 4*d* intermediate dimension, no attention bias, and QK-Norm [72, 73]:

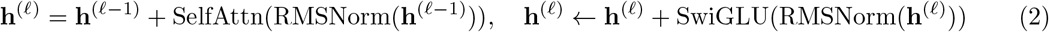

##### Cross-attention Conditioning

At designated layers (every third layer plus the final layer; 4 of 12 total in X-Cell; 11 of 34 total in X-Cell-Ultra), cross-attention blocks [74] condition the representation on external biological knowledge:

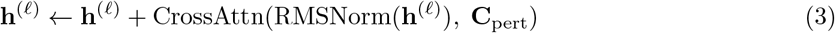

where the context **C**_pert_ ∈ ℝ^6*×d*^ consists of six prior knowledge embeddings associated with the perturbed gene. These embeddings are derived from ESM-2 protein language model [38], STRING interaction networks [41], GenePT gene representations [19], DepMap genetic dependency profiles [42], JUMP-Cell Painting morphological features [43], and the gene identity embedding itself (i.e., initialized from scGPT and updated during training in X-Cell). Because the prior knowledge embeddings have different dimensionalities, each embedding is projected to dimension *d* through a two-layer MLP with LeakyReLU activation and LayerNorm (Section A.1.2). The prior knowledge autoencoders are optimized with a reconstruction loss (Section 4.1.3). Each cross-attention block additionally includes a SwiGLU feed-forward sublayer with a Pre-LN residual connection.

##### Output Head

The CLS output **h**_CLS_ ∈ ℝ^*d*^ is expanded and concatenated with each gene’s representation, yielding [**h**_*g*_; **h**_CLS_] ∈ ℝ^2*d*^. This concatenation [**h**_*g*_; **h**_CLS_] ensures that the decoder has simultaneous access to local gene-specific features and the aggregated global cell state, allowing the model to modulate gene-level predictions based on the broader cellular context. A three-layer MLP decoder with LeakyReLU maps this to 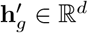. X-Cell uses a linear projection to scalar output; X-Cell-Ultra uses a tied embedding projection [75, 76]:

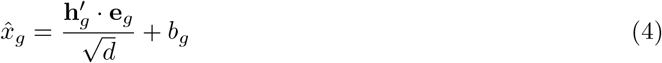

where **e**_*g*_ is the shared (un-normalized) gene embedding from the encoder, 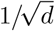 follows PaLM [77], and *b*_*g*_ is a learned per-gene bias (zero-initialized). Weight tying constrains predictions to the gene embedding space, acting as an implicit regularizer against conservative collapse [78] (Appendix A.3).

##### Training Objectives

The training loss combines seven terms, each targeting a distinct statistical property of the predicted cell distribution (all computed after CLS exclusion at position 0; Section A.2):

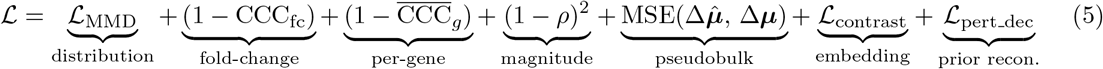

The **MMD loss** ℒ_MMD_ uses energy distance [27, 79, 80] via the `geomloss` library to match the joint distribution of *S*-cell predicted sets against ground-truth perturbed sets in *G*-dimensional gene space [81]. The **fold-change CCC** computes the concordance correlation coefficient [82, 83] between predicted and true pseudobulk fold-changes 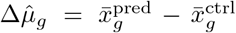 across genes, capturing both direction and magnitude (unlike scale-invariant alternatives such as Pearson *r*, which have zero gradient w.r.t. magnitude when predictions are proportional to truth). The **per-gene CCC** 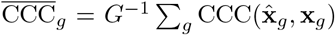 matches each gene’s marginal expression distribution across cells, preserving cell-to-cell heterogeneity that pseudobulk averaging destroys. For 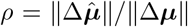, the **delta norm penalty** (1 − *ρ*)^2^ directly penalizes magnitude suppression [84]. The **contrastive loss** is InfoNCE [85] applied to CLS-derived per-cell embeddings using cosine similarity with temperature *τ* = 0.1, which controls the sharpness of the similarity distribution. Positives correspond to the centers of ground-truth embeddings for the same perturbation. The **perturbation decoder** reconstructs the five prior knowledge embeddings from the cross-attention projections via MSE, regularizing the injection of biological priors. All terms are weighted equally at 1.0; L1 output regularization is set to 0 for X-Cell-Ultra but 1.0 for X-Cell. X-Cell is trained with MMD loss, contrastive loss, and prior knowledge reconstruction while X-Cell-Ultra high-effect size curriculum training used all losses while full dataset training removed per-gene CCC.

##### Diffusion-Based Inference

At inference, predictions are refined through *T* = 4 cumulative steps [86]. At each round of prediction, 25% of the *predicted* values are incorporated into the input expression profile. Previously revealed predictions remain fixed, resulting in a cumulative update across steps. In our implementation, gene positions are assigned random ranks **r** ∼ Uniform[0, 1)^*G*^*′*. At step *t*, the model predicts the full profile from the current (partially updated) input; genes with *r*_*g*_ ≤ *α*_*t*_ (for *α*_*t*_ ∈ {0.25, 0.5, 0.75, 1.0}) that were not already revealed are accepted (clamped to ≥ 0) and scattered back into the input for the next step. The mask **M**_**pert**_ is updated correspondingly. This cumulative mode enables iterative coarse-to-fine refinement of the predictions (see Algorithm A.4 in the Appendix).

##### Curriculum Training

X-Cell-Ultra was trained in two phases following curriculum learning principles [87]. **Phase 1** (initialization): 10 epochs on the top 5% highest-effect perturbations across all seven X-Atlas/Pisces screens, using 16 nodes (128 H200 GPUs), AdamW [88] with LR = 3 × 10^−4^, cosine decay with 10% linear warmup, and optimizer step size 128 (6,144 cells per update). **Phase 2**(fine-tuning): continued training on the full dataset (100% perturbations) with 16 nodes (128 GPUs), a *fresh* AdamW optimizer (no state transfer, following [89, 90]), LR = 4 × 10^−5^ (square-root batch scaling [91]: 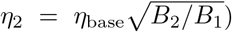, and 3% warmup. Phase 1 establishes strong perturbation signal from high-effect examples; Phase 2 extends to the full perturbation distribution at lower learning rate. Epoch 9 of Phase 1 was selected for warmstart based on held-out cell-eval metrics on iPSC and APEL datasets (Appendix A.7).

##### Test-Time Adaptation (TTA)

For zero-shot evaluation on unseen cell contexts, we optionally apply test-time adaptation [92] using control-to-control self-supervised learning. Independent pairs of non-targeting control (NTC) cell sets are drawn from the target domain, and the model is fine-tuned with MMD loss term only. Self-attention layers are adapted while cross-attention blocks are frozen (`freeze_cross_attn`) and skipped (`skip_cross_attn`) during TTA to preserve perturbation conditioning and avoid numerical instability from empty perturbation context. During test-time adaptation (TTA), optimization is performed on the output of a single-step prediction without iterative diffusion. After adaptation (∼ 200–1,024 control cell sets (*N* = 64 cells at LR ∈ {10^−6^–10^−5^}), cross-attention is re-enabled for perturbation inference. This procedure adapts the model’s internal representations to the target domain’s expression manifold without requiring any perturbation labels, analogous to reversible normalization approaches for distribution shift [93].

##### Distributed Training

All models ≥ 80M parameters were trained with mixed-precision (*bf* 16) using PyTorch FSDP2 [94] with per-layer sharding on self-attention, cross-attention, and FFN modules, following the wrapping strategy of Llama 3 [95]. Activation checkpointing is applied in the TorchTitan order [96] (checkpoint → compile → shard) to avoid dtype conflicts between FSDP2 and gradient check-pointing wrappers. Gradient clipping at norm 1.0 is applied at each optimizer step. X-Cell-Ultra achieves ∼41% MFU on 64-128 H200 GPUs.

#### 4.1.4 Benchmark Experiment Setup

To benchmark X-Cell against existing methods and baselines, we designed both **finetuning** and **zeroshot** tasks, and reported evaluation metrics implemented in the `cell-eval` package [27].

In the **finetuning** setting, we focused evaluation on a few-shot setup where perturbations are available in multiple cell types, and a proportion of perturbations in one cell type is held out for testing. While the hold-out proportion varies across **iPSC/HepG2-200, Replogle-Nadig**, and **Parse-1M** (Section 4.1.7), this generally represents a setting where some in-context perturbational data is available, but the model extends to unseen perturbations in the same cellular context. We benchmarked against **Cell2Sentence** [22], **STATE** [27], and **scGPT** [8]. In the iPSC/HepG2-200 benchmark, due to the large scale of the X-Atlas/Pisces corpus, we report a STATE model trained on a 5.5M-cell subset to better align the pre-training data with X-Cell. For Cell2Sentence, we report zero-shot readouts due to computational constraints for finetuning. X-Cell was applied directly without additional tuning. In the Replogle-Nadig and Parse-1M benchmarks, X-Cell was fine-tuned with a learning rate of 3 × 10^−4^ for 20 epochs at set size 64, using the same training objectives described in Section 4.1.3. All benchmark models were fine-tuned and evaluated using the same train–test splits as X-Cell, following the descriptions in Section 4.1.5.

In the **zero-shot** setting, we tested a more challenging setup by holding out an entire cell type (**Melanocyte Progenitors**), extrapolating to primary cells (**Human Primary T Cells**), or evaluating on distinct cell lines with chemical perturbations (**Tahoe Subset**) without observing perturbations in the test context (Section 4.1.7). In this setting, X-Cell and X-Cell-Ultra were lightly tuned on test control cells using Test Time Adaptation (TTA) (Methods 4.1.3). We report a STATE model pre-trained on the original Replogle-Nadig datasets [12][52] with full transcriptomic readout and the H1 screen from the Virtual Cell Challenge (VCC) release [55] (Section 4.1.5), representing a version of STATE trained on large-scale public data. Because the Melanocyte Progenitors test set was generated on the FLEX platform, as was the H1 screen, this combination helps align the STATE model with both the X-Cell-Ultra training corpus and the test data distribution. In the zero-shot setting, we also report two baselines, **Control Mean** and **Pert Mean**, which represent data-only predictions constructed using test control cells and log-fold changes from X-Atlas/Pisces (Section 4.1.6).

For inference and benchmarking, we used 32 fixed control cells randomly selected from NTCs in the test set to generate predictions for all test perturbations, and reported six evaluation metrics (Pearson Δ, Overlap@N, DE Direction Match, DE LogFC Spearman, MAE, and Discrimination L2 / Centroid Accuracy) from the `cell-eval` package. The same 32 control cells were used across all benchmark methods and baselines to ensure consistency.

#### 4.1.5 Benchmark Methods

##### STATE

To produce full-transcriptome predictions, we trained two versions of the STATE model [27]: a model trained on a substantial subset of four X-Atlas/Pisces screens to align with the X-Cell pre-training corpus for fair comparison, and (2) a public-data-only model trained on the original Replogle–Nadig datasets (Replogle et al.^5^ [12]; Nadig et al.^6^ [52]) together with the Virtual Cell Challenge (VCC) training release^7^ [55], referred to as Replogle+VCC.

In (1), the subset of X-Atlas/Pisces data used to train STATE consists of four cell lines (HCT, HEK, HepG2, and iPSC), with 5,367 unique perturbations and 5,545,269 cells selected to maximize validation gene coverage. In (2), the combined public Replogle+VCC datasets include five cell lines: HepG2, Jurkat, K562, RPE1, and H1. For K562, we included both the essential-genes-only and whole-genome versions. In total, the Replogle+VCC dataset contains 9,884 unique perturbations and 3,177,579 cells.

For model training, we used the SE model to generate embedding representations and the ST model to predict perturbation effects from embeddings extracted from SE. For the SE model, we downloaded the `se600m_epoch16` checkpoint from HuggingFace^8^ and followed the provided instructions to extract embedding features. To train the ST model, we adapted the official zero-shot setup released by the authors^9^ to reproduce STATE results using SE-derived embeddings.

For the Replogle–Nadig and Parse-1M benchmarks on highly variable genes, we followed the same train–test splits used for X-Cell (Section 4.1.7) and trained STATE for 20 epochs using the recommended architecture and learning rate^10,11^.

##### Cell2Sentence (C2S)

Cell2Sentence converts single-cell gene expression profiles into natural language “cell sentences”, where gene symbols are ordered by descending expression level and processed by autoregressive language models to predict perturbation responses [22]. We benchmarked the `C2S-Scale-Gemma-2-2B` model in both zero-shot and finetuning settings.

Gene expression matrices were log1p-normalized and converted into cell sentences consisting of the top 1,000 expressed genes per cell. Perturbation prediction was formulated as a sequence-to-sequence task in which the model generates a perturbed cell sentence conditioned on a control cell sentence and perturbation label. Training setup follows the C2S perturbation prediction notebook ^12^.

Due to the substantial computational cost of fine-tuning the 2B model on the X-Atlas/Pisces corpus, we report zero-shot inference using the pre-trained model on the iPSC/HepG2-200 validation set. On the Replogle-Nadig and Parse-1M benchmarks, C2S 2B model was fine-tuned on the same train-test split as X-Cell. Generated gene rankings were converted back to expression space using isotonic regression to map predicted rank positions to expression values, following the original C2S reconstruction procedure. Benchmark details are described in Appendix B.

##### scGPT

We fine-tuned the scGPT model [8] from the `scGPT_human` pre-trained weights, following the perturbation prediction architecture described in the official tutorial^13^. The model was adapted to use the MMD loss and set-based input construction, and trained at a learning rate of 5 × 10^−5^ for 10 epochs.

#### 4.1.6 Baselines

##### Pert Mean

The perturbation mean baseline assumes that the impact of a perturbation is consistent across different contexts. It estimates a perturbation effect vector Δ (in log1p CP10K gene expression space) as the average ground-truth effect across all training cell types. Let *C*_train_ denote the set of cell types observed during training, **x**_*p,c*_ the gene expression vector of a cell under perturbation *p* in cell type *c*, and ***µ***_*c*_ the mean gene expression vector of control cells for cell type *c*.

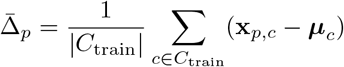

Let 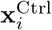 denote the gene expression vector of the *i*-th control cell in the test cell type. The predicted perturbed expression is obtained by adding the same perturbation effect vector to each control cell:

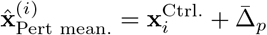

For scalability, we used 32 randomly sampled perturbed cells per perturbation and the same fixed 32 control cells used in the benchmark to estimate the perturbation effect.

##### Control Mean

The control mean baseline assumes no perturbation effect and uses the control state as the predicted perturbed state. Let 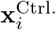 denote the gene expression vector of the *i*-th control cell (in log1p CP10K space). The prediction for each cell is therefore given by the identity mapping:

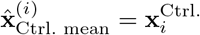

In practice, we use the same fixed set of 32 control cells employed in the benchmarking procedure as the predictions.

#### 4.1.7 Evaluation Datasets

##### iPSC/HepG2-200

In iPSC and HepG2 screens from X-Atlas/Pisces, the 200 validation perturbations were randomly selected from gene targets meeting the following criteria across all four genome-wide screens (HCT116, HEK293T, HepG2, and iPSC): (1) detectable expression in NTCs, (2) significant phenotype changes detected by the binary classifier pipeline (FDR *<* 0.05), (3) at least 100 perturbed cells per screen, and (4) ≥ 50% knockdown efficiency.

##### Replogle-Nadig

The Replogle–Nadig dataset combines Perturb-seq data from two studies across four cell lines (K562, RPE1, Jurkat, and HepG2) [12, 52]. The processed Replogle–Nadig dataset was retrieved from the release by the STATE authors [27]^14^. The processed dataset contains normalized gene expression for 643,413 cells across 2,000 highly variable genes. Following the evaluation notebook^15^, 380 HepG2 perturbations were held out as the test set, and 616,535 cells were used for training.

##### Parse-1M

We constructed the Parse-1M subset using 1,267,690 cells from donor 1 in the Parse-PBMC dataset^16^. We identified 2,000 highly variable genes from 18,308 feature space using `scanpy.pp.highly_variable_genes` with `flavor=“cell_ranger”`. To ensure compatibility with the X-Cell vocabulary, cytokine perturbations were mapped to their corresponding protein-coding gene names, and entries without valid matches were removed. After filtering, 86 cytokine perturbations with protein-coding annotations remained, comprising 1,214,863 cells. The dataset spans 18 immune cell types: CD8 Naive, CD4 Naive, CD14 Mono, NK CD56bright, Treg, CD4 Memory, MAIT, CD16 Mono, NK, B Intermediate Memory, cDC, CD8 Memory, pDC, B Naive, ILC, HSPC, NKT, and Plasmablast. Following the split 3 few-shot configuration released by the STATE authors ^17^, we held out 59 overlapping cytokines in the `CD4_Memory` cell type, and used remaining 992,862 cells for training.

##### Tahoe Subset

To test X-Cell’s zero-shot capability on chemical drug perturbation, we utilized a subset of Tahoe-100M dataset [54]. The full 100M dataset contains single-cell screening profiles of chemical drug perturbations applied to around 50 cancer cell lines, including untreated cells with DMSO as controls. We filtered the drug conditions to include all the drugs (1) having one single target MOA gene, (2) with mid-range dosage, and (3) being an inhibitor of the MOA gene ^18^. We also use the data subset of the second plate, avoiding unnecessary batch effects between plates during our zero-shot assessment. The remaining final subset has 50 cell lines and 12 drug perturbations.

##### Melanocyte Progenitors

Perturbations targeting the melanocyte cell type were held out entirely based on cell type annotations from the iPSC Multi-Diff screen (Section 4.2.12). The final gene set was derived by identifying perturbation passing a binary classifier FDR threshold of *<* 5% across four key cell populations: mesenchymal progenitors, neural crest melanocytes, epithelial ectoderm, and neuroectoderm. This core signature set of 1,317 perturbations was subsequently augmented with a curated validation set of 24 genes, including *MITF, PAX3*, and *SOX10*, selected for their established roles in neural crest specification and melanocyte differentiation. The resulting consolidated list of 1,341 genes represents a robust signature of the melanocyte progenitor developmental trajectories.

##### Human Primary T Cells

The human primary T cell dataset consists of 22 million primary human CD4^+^ T cells from four donors across three experimental conditions: resting, stimulated for 8 hours, and stimulated for 48 hours [57]. We selected donors D2 and D3 for evaluation because they exhibit a higher proportion of perturbations with phenotype changes detected by the binary classifier. For each donor, we evaluated a subset of 291 key T cell activation regulators that show differential responses across the three time points, as reported in Figure 3E of the original publication.

#### 4.1.8 Evaluation Metrics

The `cell-eval` package [27] computes per-perturbation metrics to quantify agreement between predicted and ground-truth gene expression responses. Let 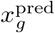 and 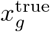 denote predicted and observed pseudobulk expression for gene *g*, and let 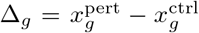 denote the perturbation effect relative to control.

**Pearson** Δ. Pearson correlation between predicted and observed perturbation effects:

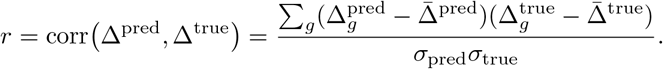

This metric measures how well the model captures the direction and relative magnitude of gene-wise expression shifts induced by a perturbation.

##### Overlap@N

Let 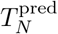 and 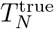 denote the sets of top-*N* ranked differentially expressed (DE) genes in prediction and ground truth. The metric is

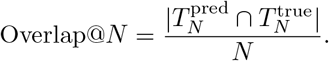

It evaluates recovery of the most strongly perturbed genes.

##### DE Direction Match

Let 𝒢_DE_ denote genes called significantly DE in both prediction and ground truth. The score is

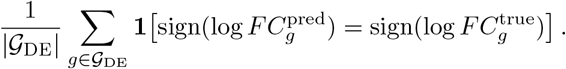

This quantifies agreement in the direction (up/down regulation) of predicted effects.

##### DE logFC Spearman

Spearman rank correlation between predicted and observed log-fold changes, restricted to genes significantly DE in the ground truth:

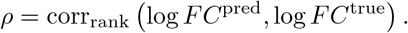

This assesses whether the model correctly ranks genes by effect size.

##### MAE

Mean absolute error between predicted and observed pseudobulk expression:

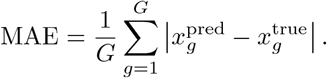

Unlike correlation metrics, MAE measures absolute deviation in expression space.

##### Discrimination L2 / Centroid Accuracy

A distance-based separation metric quantifying how well predicted perturbation signatures diverge from control signatures relative to ground truth. This metric was reported as *Centroid Accuracy* by [31]. Using Euclidean distance,

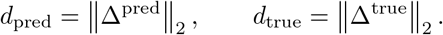

Performance reflects agreement between predicted and observed perturbation magnitudes in expression space, with stronger discrimination indicating clearer separation of perturbed from control states.

##### auROC

Area under the receiver operating characteristic curve for identifying significantly DE genes. Genes are ranked by predicted scores (e.g., − log_10_ adjusted *p*-values), and the auROC measures the probability that a randomly chosen true DE gene receives a higher score than a non-DE gene.

#### 4.1.9 Zero-shot inference of T cell inactivating perturbations

##### Inactivation Index and Distance Metrics

To quantify the phenotypic shift induced by perturbations, we projected the transcriptional profiles into a low-dimensional space using Principal Component Analysis (PCA) derived from the 2,000 most highly variable genes. We retained the top 50 principal components. Let **x**_*p*_ ∈ ℝ^50^ represent the centroid of the transcriptional profile for perturbation *p* in this PCA space. We defined ***µ***_*active*_ and ***µ***_*resting*_ as the centroids of the non-targeting controls (NTCs) in the activated and resting screens, respectively.

The distance of a perturbation to a given state was calculated using the *L*_1_ norm (Manhattan distance):

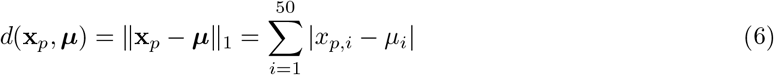

We defined the **Inactivation Index** (*I*_*p*_) as the difference between the distance to the active state and the distance to the resting state. A higher positive value indicates that the perturbation has shifted the cell state closer to the resting phenotype and further from the active phenotype:

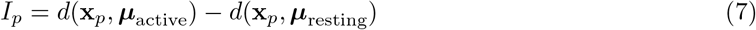

##### Baselines

We benchmarked X-Cell against a **linear baseline** to evaluate its ability to predict state changes in a zero-shot setting. This model assumes that the perturbation effect observed in the resting state (Δ_*p*_) is additive and independent of the basal cell state. Specifically, we calculated the shift (in log1p CP10k space) induced by perturbation *p* in the resting screen relative to resting non-targeting controls (NTCs) and added this shift to the activated NTC centroid:

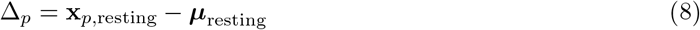

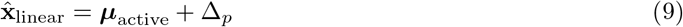

##### Statistical Analysis

We evaluated the performance on a validation set of four genes known to be critical for T cell activation as part of the CD3 complex: *CD3D, CD3E, CD3G*, and *CD247*. Significance was determined by comparing the distribution of Inactivation Indices for these candidate genes against all other perturbations using a one-sided Mann-Whitney U test. We used additional metrics from the cell-eval package such as auROC and Pearson Δ [27].

#### 4.1.10 Prior Knowledge Attribution Analysis in X-Cell

##### Cross Attention Interpretability

To characterize how individual perturbation conditions modulate attention to prior biological knowledge, we extracted cross-attention weights from the final cross-attention layer of the X-Cell model. Our approach utilized the model’s cross-attention mechanism, wherein the input gene expression sequence attends to a curated set of six prior knowledge embeddings derived from ESM-2, STRING, GenePT, DepMap, CellPainting, and scGPT. As the cross attention weights for each gene token are `softmax` normalized across these six embeddings, they provide a quantitative measure of the relative importance assigned to each knowledge source. We used the same 32 control cells from benchmark and performed inference across 380 test perturbations in the Replogle-Nadig HepG2 test set. We performed single-step prediction for simplicity, without iterative diffusion. During extraction, attention weights were averaged across heads and genes in each cell, and aggregated across cells for each perturbation.

To quantify the aggregate utilization of each knowledge source, we computed the mean cross-attention weight for each of the six embeddings across all 380 test perturbations. For Figure 3 D reporting, we removed scGPT and re-weighed the contributions. The resulting five entries were then normalized to sum to one, yielding a final distribution of attention. This distribution was visualized as a pie chart, where the proportion of each slice indicates the aggregate relative contribution of a single embedding source to the model’s cross-attention mechanism.

##### Ablation of Knowledge-Base Embeddings via Feature Permutation

To quantify the contribution of specific biological priors to model performance, we conducted a global permutation importance analysis on the five sources of knowledge-base embeddings. Each source provides a high-dimensional representation **E**_*s*_ ∈ ℝ^*G×*128^, where *G* = 19, 408 represents the set of protein-coding genes. These embeddings are integrated into the model architecture through a cross-attention mechanism, allowing the model to weigh information across different biological contexts.

###### Permutation Procedure

We evaluated the robustness of the model on the *Replogle-Nadig* HepG2 test set by systematically disrupting the information content of each embedding source *s* ∈ {1, …, 5}. For a given source, we applied a stochastic permutation function 𝒫(·) to the flattened tensor of embeddings:

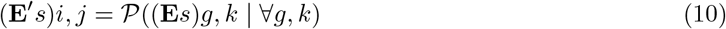

This procedure preserves the global distribution (mean, variance, and data type) of the embedding values while completely destroying the gene-specific semantic mapping and the structural relationships within the latent space. Model inference was then performed using one of the permuted embedding 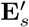 at a time while keeping all other knowledge sources and model weights fixed.

###### Quantifying Performance Degradation

The impact of each knowledge source was assessed using a suite of metrics from the `cell-eval` package, including Mean Absolute Error (MAE), L2 Discrimination Score, Pearson Δ (*ρ*_Δ_), Overlap at N, Differential Expression (DE) Direction Match, and DE LogFC Spearman correlation. To standardize the importance across heterogeneous metrics, we calculated the **Relative Performance Drop** (Δ*M*_*s*_). For metrics where higher values indicate better performance (e.g., Pearson Δ, Overlap at N), the drop was calculated as:

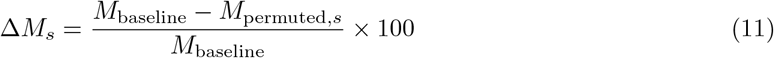

For error-based metrics where lower values are preferred (e.g., MAE), we calculated the percent increase in error:

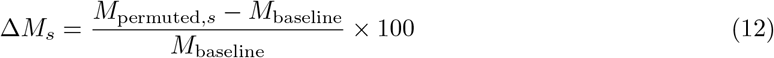

A higher Δ*M*_*s*_ indicates that the model relies more heavily on the specific biological information contained within that knowledge-base source for accurate perturbation prediction. This approach allows us to rank the five sources by their functional relevance to the HepG2 regulatory landscape and ensures that the cross-attention mechanism is effectively leveraging the provided biological priors rather than relying solely on the training distribution.

#### 4.1.11 Related Works and Extended Discussion

Early computational approaches to perturbation prediction were largely based on probabilistic latent-variable models. Conditional Perturbation Autoencoder (CPA) and its extension chemCPA model perturbational effects as structured shifts in a learned gene expression latent space, disentangling perturbations from covariates and basal cellular states [15–17]. These frameworks demonstrated that perturbation responses can be approximated as additive or compositional transformations in latent space, enabling interpolation across doses and combinations.

Subsequent work incorporated prior biological knowledge to improve extrapolation to unseen perturbations. Graph-based models such as GEARS explicitly integrate gene–gene interaction networks to enhance generalization to unseen and combinatorial perturbations [18]. Other approaches, including PRESAGE, TxPert, and GenePT, leverage pre-trained gene embeddings or structured priors derived from large biological corpora to inform perturbation modeling [19–21]. These knowledge-informed strategies provide evidence that incorporating gene regulatory or interaction structure can improve performance on unseen perturbations. However, their predictive formulation primarily captures conditional mean responses relative to control profiles and is commonly evaluated on pseudobulk summaries or deterministic reconstructions, rather than explicitly modeling the full single-cell distribution of perturbed states.

To more directly model distributional transitions, several studies have framed perturbation response as a mapping between probability distributions. Optimal transport (OT) has been used to align control and perturbed populations and to infer cellular trajectories across conditions [97]. Related neural ODE–based approaches model continuous-time transitions between cellular states [98]. In parallel, cinemaOT and related OT-based methods further formalize perturbation as a conditional distributional transport problem [25, 26]. These methods emphasize learning transformations between distributions rather than predicting point estimates, providing a principled framework for capturing heterogeneity in cellular responses. More recently, diffusion-based generative models such as scLDM and CellFlow have been proposed to model gene expression dynamics and stochastic state transitions, offering a scalable alternative for distribution-level modeling in single-cell settings [28, 29].

Large-scale pre-trained models have also been adapted to perturbation prediction. Transformer-based architectures such as scGPT and scTransformer are pre-trained on extensive public RNA-seq or single-cell corpora to learn contextualized gene representations, and subsequently fine-tuned for downstream tasks including perturbation modeling [8, 10]. Language-model–inspired approaches, such as C2S, similarly leverage large biological text or expression corpora to derive gene embeddings transferable to perturbation tasks [22]. STATE combines pre-trained representations with distributional objectives to better capture perturbation-induced variability at single-cell resolution [27]. While these efforts suggest that large-scale pre-training can improve representation quality, pre-training corpora and perturbation-specific datasets remain partially mismatched in scale and diversity, limiting systematic evaluation of scaling behavior in perturbation prediction.

### 4.2 X-Atlas/Pisces Methods

#### 4.2.1 sgRNA Library Generation, Lentiviral Production, and Single Cell Sequencing

Core methods for library construction (sgRNA design, cloning, and lentiviral packaging) and sequencing (library preparation and sequencing) were performed as previously described [39]. One modification to sgRNA library composition was that while the HCT116 and HEK293T screens targeted all protein-coding genes, subsequent screens used sgRNA libraries containing perturbations for only the genes expressed in each respective cell line as determined by bulk RNA sequencing (Section 4.2.2). Expressed genes were defined as transcripts that cumulatively account for 99% of aligned counts (Section 4.2.11).

#### 4.2.2 Bulk RNA Sequencing

100 ng of total RNA per sample was used to construct the mRNA sequencing libraries. Libraries were prepared using the Watchmaker mRNA Library Prep Kit (Watchmaker Genomics, Catalog number 7K0105-096) following the manufacturer’s protocol. xGen UDI-UMI Full-length adapters from IDT (Catalog number 10005903) were used for indexing. The final libraries were quantified using the Qubit dsDNA High Sensitivity Assay (Thermo Fisher) and their size distribution and quality were checked using the D1000 ScreenTape Analysis on the TapeStation System. Libraries were pooled in equimolar amounts. The final pool concentration was quantified via qPCR to ensure accurate loading. Pooled libraries were sequenced on a NovaSeq 6000 System (Illumina).

#### 4.2.3 iPSC Screen

ASE-9211-iPSCs (Applied Stem Cell) were engineered to express dCas9-KRAB-ZIM3 via Cas9-mediated insertion into GAPDH as previously described [99], except that the transgene insertion was targeted to the 3’ UTR of GAPDH (ASE-9211-GZ3-iPSC) just downstream of the stop codon, which was removed. The intended insertion was verified via Sanger sequencing and a normal karyotype was confirmed after editing. ASE-9211-GZ3-iPSCs were maintained in mTeSR™ Plus (Catalog number 100-0276) on Geltrex™ (Catalog number A1413202)-coated plates and tested negative for mycoplasma contamination. For the genome-scale perturb-seq experiment, 87.45 million cells were transduced in suspension at an MOI of 0.2 and then split across 33 T-150 flasks containing mTeSR Plus with 10uM Y-27632 (Catalog number 72304). Lentivirus was removed 24 hours post-transduction. Cells were then cultured in the presence of 2 µg/mL puromycin (Catalog number A1113803) for five days to enrich for transduced cells. Seven days post-transduction, cells were harvested using Accutase, then stained with Zombie NIR Fixable Viability Kit prior to bulk fixation and cryopreservation steps as described above. Aliquots of fixed cells were then thawed and sorted via FACS (Sony SH800 or BD CytoFLEX) to isolate viable (Zombie NIR-negative) cells expressing the perturbation system (dCas9-mScarlet+ and sgRNA-mNeonGreen+).

#### 4.2.4 HepG2 Screen

HepG2 cells (ATCC) were cultured in EMEM supplemented with 10 percent FBS and 100 units/mL of penicillin and 100 µg/mL of streptomycin. A CRISPRi effector consisting of dCas9-mScarlet-ZIM3 construct and a blasticidin resistance gene was stably integrated using PiggyBac transposon according to the manufacturer’s instructions (SBI). The cells were sorted to at least 95% purity for mScarlet fluorescence, expanded and cryopreserved. Cells were then thawed and expanded, and 450 million cells were seeded in 90 T150 flasks at 5 million cells per flask and transduced with the sgRNA library (MOI = 0.25). Media was exchanged the next day to remove lentivirus. Three days after transduction, cells were dissociated using Accutase and seeded in 100 T150 flasks at 15 million cells per flask in media containing 2 µg/mL. Media was refreshed with new media containing 2 µg/mL puromycin the next day, and puromycin was removed the following day with a fresh media change. Cells were harvested 7 days after sgRNA transduction with Accutase, stained with Zombie NIR Fixable Viability dye, and fixed and cryopreserved as described above. Aliquots of fixed cells were then thawed and sorted via FACS (Sony SH800 or BD CytoFLEX) to isolate viable (Zombie NIR-negative) cells expressing the perturbation system (dCas9-mScarlet+ and sgRNA-mNeonGreen+).

#### 4.2.5 Jurkat Resting and Active Screens

Jurkat (Clone E6-1) cells were cultured in RPMI 1640 supplemented with 10% FBS. The Jurkat CRISPRi line was generated by stable expression of a dCas9–mScarlet–ZIM3 CRISPRi effector carrying a blasticidin-resistance marker, delivered via PiggyBac transposition according to the manufacturer’s instructions. Cells were electroporated using the Lonza CK-116 program with a 1:2.5 ratio of transposase (SBI) to transposon, then sorted to greater than 95% mScarlet-positive purity on a Sony SH800 cell sorter. Sorted cells were maintained in medium containing 10 µg/mL blasticidin for at least 5 days prior to transduction with the genome-wide sgRNA library. For the stimulated condition, T cell activation was induced by treating cells with ImmunoCult Human CD3/CD28 T Cell Activator at a concentration of 25 µL/mL for 24 hours prior to harvest and fixation. Instead of FACS-based enrichment, magnetic-activated cell sorting (MACS) was used to minimize shear stress on the fixed lymphocytes. Thawed cells were processed using the Dead Cell Removal Kit (Miltenyi Biotec, Catalog number 130-090-101) to deplete lysed cells. Wide-bore pipette tips were used throughout the thawing and enrichment process to further reduce physical stress on the cells.

#### 4.2.6 iPSC Multi-Diff Screen

Approximately 1.27e9 ASE-9211-GZ3-iPSCs (Section 4.2.3) were transduced in suspension at an MOI of 0.2 and then split across 41 T-150 flasks containing mTeSR™ Plus with 10uM Y-27632. Lentivirus was removed 24 hours post-transduction. Two days post-transduction, iPSCs were harvested into a single cell suspension using Accutase. The MOI was determined via flow cytometry and the cells were subsequently seeded into 38 6-well plates at a density of 10.9e6 cells per well in the presence of 2ug/mL puromycin. 24 hours post-seeding (D0 differentiation), the media was replaced with STEMdiff™ APEL™2 Medium (STEMCELL Technologies, Catalog number 05275). Media changes were performed on D1-3, D5-8, and D9-12. No media change was performed on D4. The cells were harvested on D13 using Accutase, then stained with Zombie NIR Fixable Viability Kit prior to bulk fixation and long-term storage at -80C using 10X Genomics Fixed RNA Flex assay reagents. Aliquots of fixed cells were then thawed, sorted via FACS (Sony SH800 or BD CytoFLEX) to isolate viable (Zombie NIR-negative) cells expressing the perturbation system (dCas9-mScarlet+ and sgRNA-mNeonGreen+), and prepared for long-term storage at -80C. All cell sorting occurred within three weeks of fixation.

#### 4.2.7 Optimized FiCS-Perturb Workflow Modifications

To enable genome-scale throughput while mitigating batch effects and lysis-induced sequencing noise, we implemented an optimized “Fix-Freeze-Enrich” workflow that modifies the original FiCS-Perturb protocol. Unlike the previous method where cells were sorted immediately after fixation, this workflow decouples fixation from library preparation to allow for bulk banking. Cells were harvested and fixed with 1 mg/mL dithiobis (succinimidyl propionate) (DSP) for 30 minutes at room temperature. Following quenching and washing, cells were immediately resuspended in Bambanker™ freezing media supplemented with 0.2 U/µL RNasin Plus RNase Inhibitor and frozen in bulk aliquots of 10–50 million cells at - 80°C. On the day of library preparation, bulk aliquots were thawed rapidly at 37°C until a small ice crystal remained. To prevent osmotic shock and cell lysis, thawed cells were diluted slowly with cold 10X Cell Buffer (DPBS + BSA), adding 1 mL at a time followed by gentle inversion before bringing the final volume up to 15–50 mL. Thawed cells were then enriched to remove non-viable cells and debris accumulated during the freeze-thaw process immediately prior to quantification and “superloading” onto the 10x Genomics Chromium GEM-X 5’ Chip at a target density of 100,000 cells per lane.

#### 4.2.8 10X Flex-Based Single Cell Transcriptome Profiling and sgRNA Detection (Flex Perturb-seq)

For the iPSC Multi-Diff screen, aliquots of fixed and sorted cells were thawed and used as input for the Flex Fixed RNA Assay (v1) with custom probes spiked-in to detect the CRISPRi and sgRNA constructs and recover the identity of sgRNA perturbations. Six sets of 16 hybridization reactions were performed. Individual hybridizations were then pooled and washed, followed by GEM generation (34 total GEMs) aiming to recover 320,000 cells per GEM lane. Sequencing libraries were generated according to the manufacturer’s instructions with modifications made to the hybridizations, the Pre-Amplification PCR step, and the Feature PCR, according to the Technical Guidance document “CRISPR Screening with GEM-X Flex Gene Expression” (CG000814 Rev A). Sequencing was performed using an Illumina 25B NovaSeq X kit, with three Gene Expression libraries and three paired CRISPR libraries pooled per run. Each pool was sequenced twice, targeting a recovery of 320,000 cells per lane. Gene expression libraries were sequenced to a depth of 50,000 reads per cell, while CRISPR libraries targeted 5,000 reads per cell per sample. The final Gene Expression and CRISPR libraries were separately pooled to 10 nM, determined via high-sensitivity DNA Qubit quantification and TapeStation D1000 ScreenTape reagents using the average fragment size range of 200 bp to 1000 bp. qPCR was performed with the 10nM final libraries to verify the final concentration prior to sequencing using the CFX Opus 384 Real-Time PCR System (Catalog number 12011452) with the Kapa Library Quantification Kit Universal qPCR Master Mix (Catalog number KK4824). Following qPCR, the 10 nM pools were diluted with Qiagen EB buffer to adjust the final concentration to 2 nM with a 10% PhiX spike-in. The final 2 nM pooled libraries were then denatured and sequenced following the Illumina Novaseq X protocol, with a final loading concentration target of 170 pM. Sequencing parameters were set to 28/10/10/90.

#### 4.2.9 Basecalling, Alignment, and sgRNA Calling

bcl2fastq2 (v2.2) was used for basecalling and FASTQ demultiplexing. The following cellranger versions were used for alignment, cell barcode calling, and UMI counting: v8.0.0 (HCT116), v8.0.1 (HCT116, HEK293T, iPSC, HepG2), v9.0.1 (Jurkat Resting, Jurkat Active), and v10.0.0 (iPSC Multi-Diff). The 10x Genomics GRCh38 2024-A pre-built genome was used as the reference transcriptome for the HCT116, HEK293T, HepG2, iPSC, Jurkat Resting, and Jurkat Active screens. A custom reference genome (GRCh38 2024-A) and probe set (Human Transcriptome v1.1.0) with BFP, mScarlet, and dCas9 added were used for aligning the iPSC Multi-Diff screen. After alignment, QC metrics, such as mean reads per cell, median genes per cell, and median CRISPR UMIs per cells, were examined for each GEM group to ensure sufficient sequencing depth and coverage.

sgRNA calling was performed as previously described [39], with an additional optimization step to determine the minimum UMI threshold. Different thresholds ranging from 3 to 50 were evaluated, and the threshold that maximized the number of valid two-guide cells (cells assigned exactly one matched guide pair) was selected. Guides with UMI counts below the selected threshold were removed, and each cell was then assigned zero, one, or multiple sgRNA identities based on the remaining guide calls. Only cells with the expected two-sgRNA same-gene-target pair were used for subsequent downstream analyses.

#### 4.2.10 QC Metrics for X-Atlas/Pisces

The following metrics were used to assess the quality of X-Atlas/Pisces. Mean reads per cell: The total number of gene expression reads divided by the number of cell-associated barcodes. Median UMIs per cell: The median number of UMI counts per cell-associated barcode. Median genes per cell: The median number of genes detected (≥ 1 UMI) per cell-associated barcode. Mean CRISPR reads per cell: The total number of CRISPR reads divided by the number of cell-associated barcodes. Median CRISPR UMIs per cell: The median number of CRISPR UMIs per cell-associated barcode (summed over all recognized guide sequences). # perturbations in screen: The total count of unique perturbations for which at least one valid, dual-guide cell was identified. Median number cells per perturbation: The median cell count per perturbation calculated from the population of cells assigned a valid, dual-guide pair. Median on-target KD % per perturbation: The KD % for each perturbation was calculated by comparing the mean normalized expression of the target gene in the perturbed population (*Ē*_*target*_) to its expression in the NTC population (*Ē* _*NTC*_) using the formula: 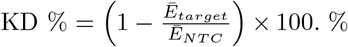 of perturbations with significant phenotype: % of perturbations with FDR *<* 0.05 as determined by the binary classifier (Section 4.2.13).

#### 4.2.11 Defining Genome-Scale Libraries for X-Atlas/Pisces

Bulk RNA-seq (Section 4.2.2) was used to determine expressed genes for the HCT116, HEK293T, HepG2, iPSC, Jurkat Resting, and Jurkat Active screens. Reads were demultiplexed and gene counts and TPM were calculated using STAR (v2.7.10a) -Salmon (v1.10.1) as implemented in the Nextflow nf-core pipeline (v3.16.0) and aggregated by ENSEMBL gene ID. Gene expression level was calculated as transcripts per million (TPM). Expressed genes were defined as genes that collectively account for 99% of the TPM counts.

For the Jurkat Resting and Active screens, 20 samples across 5 conditions (resting, ImmunoCult, Transact, CD3/CD28 Dynabeads, and PMA/Ionomycin; 4 replicates each condition) were demutiplexed and aligned using the same parameters described above. Gene expression level was assessed as transcripts per million (TPM), with expression variability scored as the standard deviation of the log10(TPM + 1), with the top 500 considered highly variable. To systematically select active genes in each sample, we first ranked genes by TPM from highest to lowest and then calculated the cumulative fraction of total TPM over that list. At a given TPM threshold (e.g. 99%), active genes were those that accounted for that much cumulative fraction of TPM. Genes found to be active in two or more samples from a single stimulation condition were marked as active in that condition. Differentially expressed genes between resting and active Jurkat cells were identified using PyDESeq2 (v0.5.3). Low expression genes were filtered prior to testing. Only genes that had a count of *>*10 in *>*4 samples were included. For each of the four stimulation conditions, gene expression levels were contrasted with the baseline of resting using a Wald test. Within each contrast, *p*-values were adjusted for multiple testing using Benjamini-Hochberg correction, with an FDR of *<* 0.05 considered significant. Significant genes from any stimulation condition were included in the final library.

For the iPSC Multi-Diff screen, the genome-scale library was determined from a combination of bulk and single-cell RNA-seq. Using single-cell RNA-seq, a multi-timepoint iPSC Multi-Diff screen was performed and collected at days 2, 6, 9, and 13. Cells were annotated into their respective cell types and pseudobulk expression profiles were generated per cell type. Expressed gene lists were generated from bulk and per cell type pseudobulks separately by retaining all genes that contributed to the top 99% of expressed transcripts. The final genome-scale library was defined as the union of the expressed genes defined from the bulk RNA-seq profile and from the pseudobulked per cell type profiles.

#### 4.2.12 iPSC Multi-Diff Cell Type Annotation

Cell x gene count matrices were processed using scanpy (v1.10.1). Cells were filtered based on the following criteria: total counts (5000 *<* n counts *<* 130,000); total genes (1,000 *<* n genes *<* 11,000); percent mitochondrial reads (pct mt ≤ 10); contain valid 2-guide pair. Genes detected in ≥ 3 cells were retained for downstream analysis using scanpy.pp.filter genes (min cells=3). Counts were normalized to counts per 10K and log2-transformed. Highly variable genes were detected using scanpy.pp.highly variable genes(n top genes=3000, n bins=20, flavor=“seurat”). Dimensionality reduction with PCA was computed using scanpy.pp.pca(n comps=50, svd solver=“arpack”). Nearest neighbors was calculated using scanpy.pp.neighbors() using the default parameters. Clustering was calculated using scanpy.tl.leiden(resolution=0.50, n iterations=5, flavor=“igraph”). Marker genes were identified per cluster using scanpy.tl.rank genes groups(n genes=50, method=“wilcoxon”). A reference dataset was created with these processing steps and manually annotated using de novo derived marker genes per cluster in addition to curated marker gene sets for germinal layers. All subsequent count matrices from individual sequencing batches were annotated to the reference object using scanpy.tl.ingest() with default parameters.

#### 4.2.13 Binary Classification

All screens in X-Atlas/Pisces were evaluated using a binary classifier as previously described [39], with two additional modifications: (1) perturbations with at least 10 cells were processed and (2) multiple testing correction was applied using the Benjamini-Hochberg procedure. Due to the variable number of cells recovered across the annotated cell types in the iPSC Multi-Differentiation screen, we report the binary classifier results obtained from analyzing all cells in the dataset without subsetting by cell type.

#### 4.2.14 Recall of Known Biological Networks

Biological coverage across X-Atlas/Pisces was assessed using the EFAAR framework [100]. Perturbations with *<*10 cells were excluded, while remaining perturbations and non-targeting controls were downsampled to a maximum of 100 cells per screen and 100 cells per GEM, respectively. Transcriptomic profiles were embedded via PCA, outlier-filtered, and cross-batch aligned using Typical Variation Normalization (TVN). After mean-aggregating the aligned profiles by perturbation, pairwise gene relationships were inferred using cosine similarity and converted to relative ranks within each experiment.

Gene similarity ranks were compared against reference interactions from CORUM [101] and STRING [102] (v12.0, score ≥ 950). Recall was computed using gene pairs in the top and bottom 5% of similarity ranks, including only pairs present in both the dataset and reference database. To assess the impact of perturbation pairs across combined contexts, cumulative recall metric was calculated across all possible combinations the screens (reporting mean and standard deviation). The union of top and bottom 5% gene pairs was calculated, retaining the most extreme rank for pairs detected in multiple screens. To evaluate cumulative performance across one to seven screens, recall was calculated for every possible subset. For each subset size (e.g., all 21 possible combinations for a two screen subset), we report the mean and standard deviation of the recall.

#### 4.2.15 Identification of Context-Independent and Context-Dependent Perturbations

Binary classifier F1 scores were compiled across seven screens (HCT116, HEK293T, HepG2, iPSC, Jurkat Resting, Jurkat Active, and iPSC Multi-Diff) for perturbations with a maximum F1 score ≥ 0.75 and present in ≥ 3 screens. The resulting F1 score matrix was clustered into 10 groups using k-means clustering (random seed=42), with missing values filled by 0 for distance calculations. Clusters were ordered by size in descending order, and genes within each cluster were sorted by mean F1 score across screens. Columns (screens) were hierarchically clustered using average linkage with Euclidean distance on column-mean-imputed data. The heatmap was rendered using seaborn.cluster() (seaborn v0.13.2), with cluster assignments annotated as row colors. To identify context-independent and context-dependent perturbations, a specificity score was computed for each cluster as the mean F1 in the screen of interest minus the mean F1 across all other screens. Gene set enrichment analysis (GSEA) was performed on genes from the context-independent cluster (highest mean F1 across all screens) and the most screen-specific clusters using gseapy.enrich() (gseapy, v1.1.4) against MSigDB v2025.1 gene sets (WikiPathways, Reactome, GO Biological Process, GO Cellular Component, GO Molecular Function, and Hallmark), with the full set of heatmap genes as background and FDR *<* 0.05.

#### 4.2.16 Visualization of Perturbations by Screen on UMAP

Features (genes) derived from the binary classifier across all seven screens (n=39,447 perturbations) were aggregated into a combined data matrix. Perturbations were excluded from the embedding if their *p*-value from the binary classifier was ≥ 0.05 or if the only detected feature was the perturbed gene itself. Feature gene lists were further restricted to a set of 19,082 protein-coding genes, defined by the intersection of GENCODE annotations and the ESM-2 vocabulary. This filtering yielded a final analytical set of 35,016 perturbations.

For each perturbation, detected feature genes were encoded into a sparse binary vector using sklearn.preprocessing.MultiLabelBinarizer (scikit-learn v1.7.2). The resulting sparse matrix (35,016 perturbations × 11,351 genes) was transformed applying L2-normalized term frequency-inverse document frequency (TF-IDF) weighting via sklearn.feature extraction.text.TfidfTransformer(norm=“l2”, sublinear tf=False). To generate a dense embedding, dimensionality reduction was performed using sklearn.decomposition.TruncatedSVD(n components=100) to retain the top 100 singular vectors. The SVD embedding was visualized using umap.UMAP(n neighbors=30, min dist=0.005, metric=“cosine”, n components=2) (umap-learn v0.5.9). Density-based clustering was subsequently performed on the UMAP embeddings using sklearn.cluster.HDBSCAN(min cluster size=20, min samples=5, cluster selection method=“eom”, metric=“euclidean”). This assigned 88.1% of the perturbations into 253 distinct clusters, leaving the remaining 11.9% as unassigned noise.

To assess functional coherence, the unique set of perturbed genes within each cluster was tested for enrichment against the CORUM database of human protein complexes using gseapy.enrich() (gseapy v1.1.11). Following Benjamini-Hochberg correction, 223 of the 253 clusters exhibited significant enrichment for at least one CORUM complex (FDR *<* 0.05). Protein-protein interaction (PPI) enrichment was evaluated using the STRING database (network v11.5). For each cluster, an observed score was calculated as the mean STRING combined score among all intra-cluster gene pairs. A matched background null distribution was generated by averaging intra-cluster PPI scores across 200 random cluster label permutations using numpy.random.default rng(42).permutation (numpy v1.26.4), preserving original cluster sizes. Observed cluster scores were significantly higher than their corresponding null scores (paired t-test using scipy.stats.ttest rel (scipy v1.16.3); n=253 clusters, *t*=8.60, *p*-value=8.44 × 10^−16^).

For the visualization of specific macromolecular complexes, spatially coherent clusters were manually curated for strong CORUM enrichments. The union of genes from the selected clusters was mapped to corresponding structural models (PDB: 3J9M for the mitoribosome [47]; PDB: 5XTD for Complex I [48]). Structures were rendered using the pymol2 API. Chains corresponding to detected genes were visualized using cmd.fetch() and cmd.show(“cartoon”) and were mapped to a shuffled cyan-to-lime-green color gradient. Undetected chains were colored medium gray to provide visual contrast.

## 5 Data and Model/Code Availability

### 5.1 Data Availability

The following data will be uploaded to https://huggingface.co/datasets/Xaira-Therapeutics/ X-Atlas-Pisces: (1) all cells from X-Atlas/Orion (cells with a valid guide pair from HCT116 and HEK293T screens), (2) cells containing the 200-gene test perturbation set from the HepG2 and iPSC screens, and all cells with a valid guide pair from the Jurkat Resting and Jurkat Active screens.

### 5.2 Model and Code Availability

To facilitate the reproducibility of our results and encourage community adoption of the *X-Cell* architecture, we have made X-Cell (55M) model artifacts publicly available. The trained model weights are hosted on the Hugging Face Model Hub at https://huggingface.com/Xaira-Therapeutics/X-Cell. Furthermore, we provide software and tutorials on GitHub at https://github.com/xaira-therapeutics/x-cell.

The dataset and trained model weights are released under the CC BY-NC-SA 4.0 license.

## 6 Declarations

All authors are employees of Xaira Therapeutics and may hold company equity.

## Appendix A X-Cell Model Specification

### A.1 Architecture Details

#### A.1.1 Input Encoding

Each training sample is a tuple (**X**_ctrl_, **X**_target_, **T, M**_pert_, *p*) where **X**_ctrl_, **X**_target_ ∈ ℝ^*S×G*^*′* are control and perturbed expression matrices over *S* cells and *G*^*′*^ subsampled genes, **T** ∈ ℤ^*S×G*^*′* contains gene vocabulary indices, **M**_pert_ ∈ {0, 1}^*S×G*^*′* is a binary perturbation mask indicating which gene positions contain revealed perturbed signal, and *p* identifies the perturbation. The batch is reshaped from (*B, S, G*^*′*^) to (*B* · *S, G*^*′*^) for token-level processing.

During training, the observed expression input 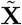 is constructed by mixing control and perturbed values according to the perturbation mask:

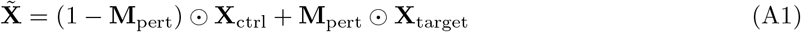

where ⊙ denotes elementwise multiplication.

Three encoders produce *d*-dimensional representations:

##### Gene identity encoder

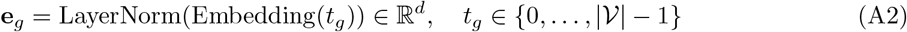

where 𝒱 is the gene vocabulary containing |𝒱| ≈ 19,400 protein-coding genes plus special tokens (`<cls>`, `<mask>`) and perturbation gene aliases.

##### Continuous value encoder

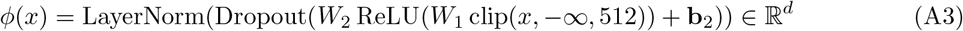

with *W*_1_ ∈ ℝ^*d×*1^, *W*_2_ ∈ ℝ^*d×d*^.

##### Perturbation mask encoder

The binary perturbation mask is embedded through a learned embedding table followed by LayerNorm:

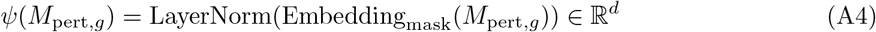

which indicates whether the input value at position *g* corresponds to control or revealed perturbed signal.

##### CLS token (learned)

The CLS token at position 0 is a learned embedding **c** = Embedding_CLS_(0) ∈ ℝ^*d*^, independent of gene, value, and mask encoders. Gene positions 1 through *G*^*′*^ are encoded normally, and the CLS embedding is prepended:

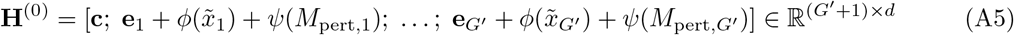

#### A.1.2 Perturbation Context Encoding

For each perturbation *p*, six prior knowledge embeddings are projected to dimension *d*:

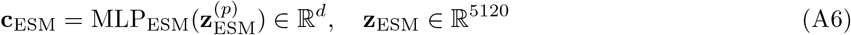

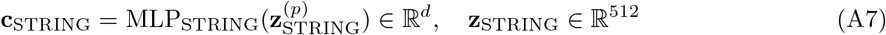

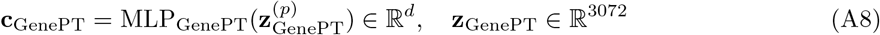

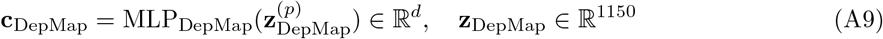

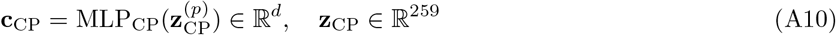

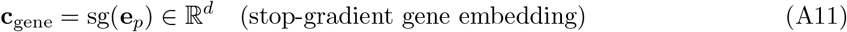

where each 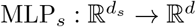 is a two-layer network with LeakyReLU activation and LayerNorm output. The six tokens form the cross-attention context **C**_pert_ = [**c**_ESM_; **c**_STRING_; **c**_GenePT_; **c**_DepMap_; **c**_CP_; **c**_gene_] ∈ ℝ^6*×d*^, replicated across the *S* cells in each set.

Missing embeddings (e.g., when a gene lacks a STRING entry) are zero-imputed and flagged via a boolean mask **m**_ext_ ∈ {0, 1}^6^ passed as `key_padding_mask` to cross-attention.

#### A.1.3 Transformer Encoder

The encoder consists of *L* self-attention layers with interleaved cross-attention at designated indices ℐ_cross_ ⊂ {0, …, *L* − 1}.

##### Self-attention with QK-Norm

For layer *ℓ*:

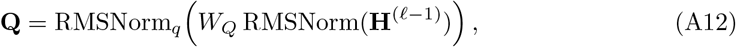

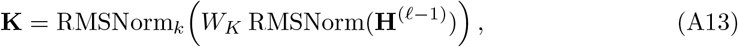

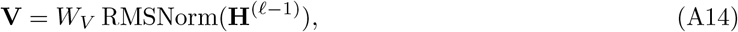

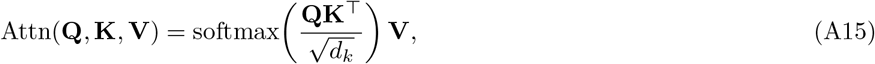

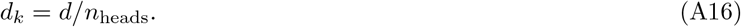

where RMSNorm_*q*_, RMSNorm_*k*_ are per-head RMSNorm (learnable scale, no bias) applied independently to each attention head [72]. All projection matrices *W*_*Q*_, *W*_*K*_, *W*_*V*_, *W*_*O*_ ∈ ℝ^*d×d*^ omit bias terms (LLaMA convention).

##### RMSNorm

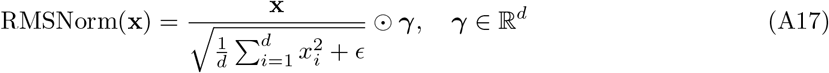

##### SwiGLU feed-forward network

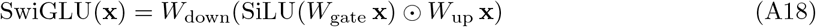

with 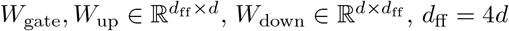, all without bias.

##### Pre-LN residual block

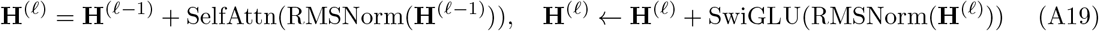

##### Cross-attention block

At layers *ℓ* ∈ ℐ_cross_, a cross-attention block is applied after the self-attention residual:

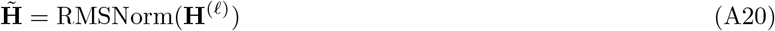

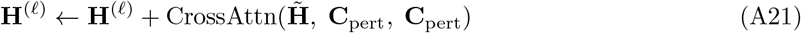

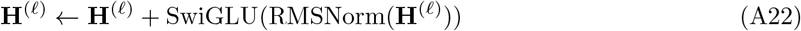

where the query is derived from 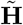 (gene representations) and the key/value from **C**_pert_ (perturbation context). This conditions each gene representation on biological prior knowledge about the perturbation. A final RMSNorm is applied after all *L* layers.

To facilitate gradient flow and preserve the initial identity of gene signals, a global residual connection is optionally applied:

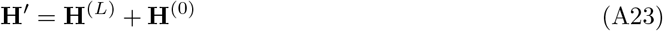

where **H**^(0)^ is the initial input representation before the transformer blocks.

#### A.1.4 Cell Embedding and Decoder

The CLS token output 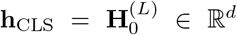 serves as a cell-level embedding. It is expanded and concatenated with gene-level outputs:

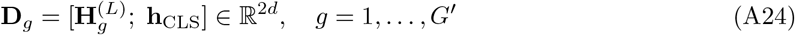

The expression decoder is a three-layer MLP with LeakyReLU (*α* = 0.01):

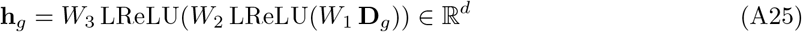

##### Tied embedding output

The predicted expression for gene *g* is computed via Eq. 4:

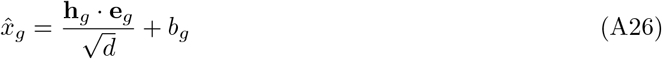

where **e**_*g*_ are the raw (un-normalized) gene embeddings shared with the encoder, and *b*_*g*_ ∈ ℝ is a learned per-gene bias via `nn.Embedding`. The 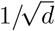 scaling prevents logit explosion at large hidden dimensions [77].

#### A.1.5 Perturbation Decoder (Auxiliary)

To regularize cross-attention, the model reconstructs each prior knowledge embedding from the cross-attention output via per-source MLPs:

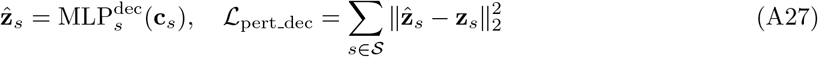

where 𝒮 = {ESM, STRING, GenePT, DepMap, CP}.

### A.2 Training Objectives

All loss terms operate after CLS exclusion: 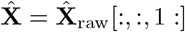, removing position 0.

#### A.2.1 MMD Loss (Energy Distance)

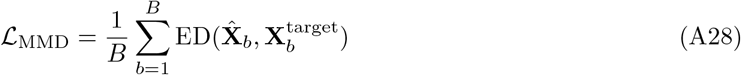

where ED is the energy distance [27, 79] computed via the `geomloss` library with `kernel=“energy”`, `blur=0.05`, `scaling=0.5` [81]. Each 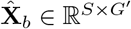 is a set of *S* predicted cells, compared against *S* ground-truth perturbed cells. The energy distance captures distributional properties (variance, multi-modality) beyond what MSE provides.

#### A.2.2 Fold-Change CCC Loss

The concordance correlation coefficient (CCC) [82] measures agreement between predicted and true pseudobulk fold-changes:

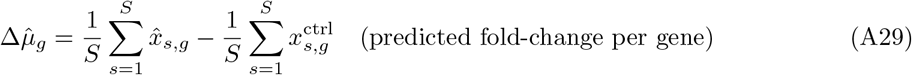

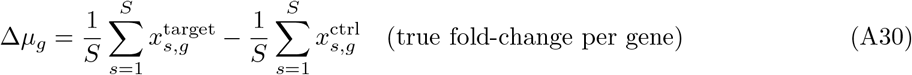

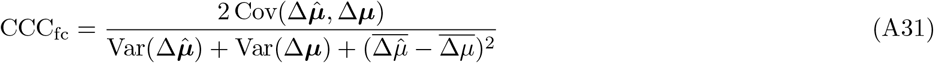

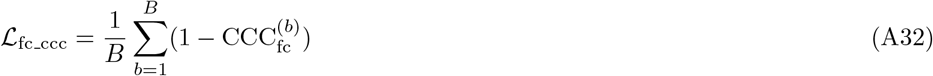

where statistics are computed across genes within each batch sample. Unlike scale-invariant losses (Pearson, cosine), CCC penalizes both directional and magnitude mismatch, which ablation studies showed were critical for combating conservative collapse (see Table A1).

**Table A1.** Architecture and loss ablation (1.75B model, Replogle–Nadig, 6 epochs, loss = 64). Each row changes one factor from the row above.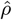: *δ*-norm ratio (ideal = 1); CA: centroid accuracy.

| Configuration | MAE↓ | DE-$r$↑ | $\hat{\rho}$ | CA↑ |
| --- | --- | --- | --- | --- |
| MMD + Pearson + cosine | 0.176 | 0.050 | 0.22 | 0.81 |
| + CCC, δ-norm penalty | 0.178 | 0.123 | 0.32 | 0.83 |
| + remove hidden residual | **0.163** | 0.150 | 0.43 | 0.84 |
| + tied embeddings, CLS fix | 0.172 | **0.327** | **0.51** | **0.86** |

#### A.2.3 Per-Gene CCC Loss

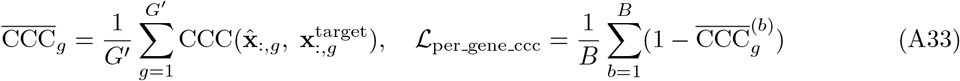

where CCC is computed across cells (dim *S*) for each gene independently. This matches per-gene marginal distributions, complementing MMD’s joint distribution matching.

#### A.2.4 Delta Norm Penalty

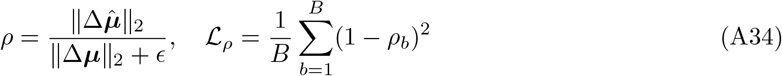

A direct anti-collapse regularizer: *ρ* ≪ 1 indicates the model suppresses perturbation magnitude (predicting expression close to control). The squared penalty is symmetric around *ρ* = 1.

#### A.2.5 Delta Pseudobulk MSE

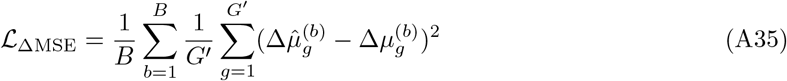

Per-gene L2 error on fold-changes. Provides absolute magnitude signal complementing the correlation-based CCC.

#### A.2.6 Contrastive Embedding Loss

InfoNCE [85] on CLS-derived cell embeddings:

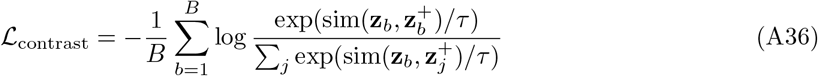

where **z**_*b*_ is the predicted cell embedding, 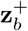 is the center of the ground-truth embedding for the same perturbation, negatives are centers of other perturbations, sim is cosine similarity, and *τ* = 0.1.

#### A.2.7 Complete Objective

All seven terms are summed with equal weight *w* = 1.0 (L1 regularization weight set to 0):

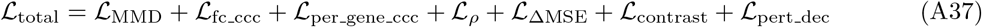

##### Gradient RMS monitoring

To diagnose gradient balance, per-loss gradient 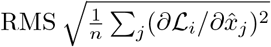 is logged for the first 100 training steps. This revealed that scale-invariant losses (Pearson, cosine) diluted the MMD gradient without fixing collapse, motivating the switch to CCC.

### A.3 Architecture and Loss Ablations

Large models (≥ 1.75B parameters) exhibited conservative collapse [84]: correct perturbation direction (DE Pearson *r >* 0) but suppressed magnitude (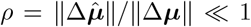, *ρ* ≈ 0.17–0.22 for early 3.5B runs). A systematic ablation on the 1.75B model (Replogle-Nadig, 6 epochs) isolated five root causes (Table A1; Supplementary Figure 7A):

**1. Post-LN** → **Pre-LN**. Post-LN collapsed at *L >* 12 (*ρ <* 0.1). Pre-LN [69] normalizes before attention, stabilizing deep training.

**2. Pearson/cosine** → **CCC**. Scale-invariant losses have 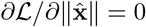 when 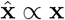. CCC’s (*µ*_*x*_ − *µ*_*y*_)^2^ denominator term provides non-zero magnitude gradient [82]. Gradient RMS: Pearson *<*5% of total vs. CCC 20–40%.

**3. Residual removal**. The skip **H**^(*L*)^ += **H**^(0)^ creates a lazy shortcut [103–105]. Removing it: MAE 0.178 → 0.163.

**4. L1 regularization**. 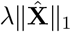 directly penalizes magnitude; set *λ* = 0.

**5. Tied embeddings**. Weight tying [75] with 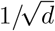 scaling [77] constrains outputs to the gene embedding space, preventing neural regression collapse [78]: *ρ*: 0.43 → 0.51, DE-*r*: 0.150 → 0.327.

**Table A2.** X-Cell model configurations.

| Parameter | X-Cell (55M) | X-Cell-Ultra (4.87B) |
| --- | --- | --- |
| Layers $L$ | 12 | 34 |
| Hidden dim $d$ | 512 | 2560 |
| Attention heads | 8 | 40 |
| Head dim $d_k$ | 64 | 64 |
| FFN dim $d_{\text{ff}}$ | 512 ($1 \times d$) | 10240 ($4 \times d$) |
| FFN activation | ReLU | SwiGLU |
| Normalization | Post-LN (LayerNorm) | Pre-LN (RMSNorm) |
| Cross-attn layers | 4 (indices 2,5,8,11) | 11 (every 3rd + final) |
| QK-Norm | No | Yes |
| Attention bias | Yes | No |
| Output head | MLP | Tied embeddings |
| Hidden residual | No | No |
| Pre-trained init | scGPT | From scratch |
| Gradient checkpointing | No | Yes |

**Table A3.** Training hyperparameters for Phase 1 and Phase 2 curriculum.

| Parameter | Mini (60M) | X-Cell-Ultra Phase 1 | X-Cell Ultra Phase 2 |
| --- | --- | --- | --- |
| Dataset | Replogle-Nadig | Top 5% high-effect | 100% all perturbations |
| Nodes (GPUs) | 1 (8) | 12 (96) | 16 (128) |
| Set size $S$ | 64 | 64 | 64 |
| Optimizer step size | 64 | 96 | 128 |
| Effective batch (cells) | 4,096 | 6,144 | 8,192 |
| Epochs | 6 | 10 | 10 |
| Optimizer | AdamW | AdamW | AdamW (fresh) |
| $\beta_1, \beta_2$ | 0.9, 0.999 | 0.9, 0.999 | 0.9, 0.999 |
| Peak LR | $10^{-4}$ | $3 \times 10^{-4}$ | $4 \times 10^{-5}$ |
| Weight decay | 0.01 | 0 | 0 |
| LR schedule | Cosine + warmup | Cosine + warmup | Cosine + warmup |
| Warmup | 10% | 10% | 3% |
| Grad clip norm | 1.0 | 1.0 | 1.0 |
| Context length $G'$ | Full ($\sim 19\text{k}$) | 4,000 | 4,000 |
| Mixed precision | bf16 | bf16 | bf16 |
| FSDP strategy | N/A | Per-layer | Per-layer |
| Warmstart | scGPT embeddings | — | Phase 1 epoch 9 |

#### Training efficiency

Per-layer FSDP2 wrapping [94] reduced step time from ∼17 s to ∼12.3 s (28%). X-Cell-Ultra achieves 41% MFU on 64 H200 GPUs, comparable to Llama 3’s 38–43% [95] and *>*20× above prior single-cell models (∼1–16%; Supplementary Figure 7B).

#### QK-Norm

Per-head RMSNorm on Q, K [72] prevents bf16 attention logit explosion at 34 layers. Modest metric improvement; primarily a stability measure for production training.

### A.4 Diffusion Mask Mechanism

#### Training

For each training sample, a control mask strategy *α* is drawn uniformly from {0.25, 0.5, 0.75, 1.0}. The last ⌊*α* ·*G*^*′*^ ⌋ gene positions (in a fixed random permutation) have their control expression replaced with target expression:

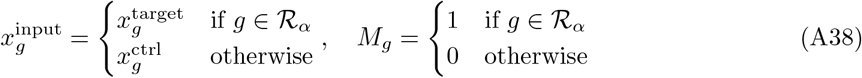

where ℛ_*α*_ is the set of revealed positions. The model is trained to predict the full **X**_target_ from this partially revealed input.

#### Inference (cumulative mode)

At inference, predictions are iteratively refined across *T* = 4 steps:

1. Initialize: **X**^(0)^ = **X**_ctrl_, **M**^(0)^ = **0**, assign random ranks **r** ∈ [0, 1)^*G*^*′* to genes.
2. For *t* = 1, …, *T* with reveal fractions *α*_*t*_ ∈ {0.25, 0.5, 0.75, 1.0}:
  a. Forward pass: 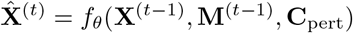
  b. Select newly revealed genes: 𝒩_*t*_ = {*g*: *r*_*g*_ ≤ *α*_*t*_} \ {*g*: *r*_*g*_ ≤ *α*_*t*−1_}
  c. Update: 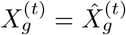 for 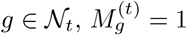 for *g* ∈ 𝒩*t*
3. Return 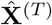 as the final prediction.

### A.5 Test-Time Adaptation Details

Test-time adaptation (TTA) enables zero-shot generalization to unseen cell contexts by adapting the model’s self-attention representations to the target domain’s expression manifold, without requiring perturbation labels [92].

#### Data construction

In control-to-control (ctrl2ctrl) mode, the dataloader samples independent pairs of NTC cell sets from the target domain. Each training pair consists of two independently drawn NTC sets of *S* cells, yielding self-supervised distribution-matching signal. The model learns the target domain’s expression statistics (gene-gene correlations, variance structure, cell-type-specific patterns) without any perturbation information. This is implemented via `ctrl_to_ctrl=True` in the dataloader configuration, with `ctrl_mask_strategy=“none”` to ensure **M**_pert_ = **0** (no perturbation mask signal).

#### Model configuration

Two flags jointly control the adaptation:

- `skip_cross_attn=True`: passes `label_ctx=None` to the transformer, bypassing all 11 cross-attention blocks. This avoids numerical instability from all-masked attention keys (no meaningful perturbation context exists for NTC pairs, since all external embedding masks would be active).
- `freeze cross attn=True`: cross-attention block parameters (∼ 1.15B on X-Cell-Ultra) have `requires_grad=False`. After TTA, `skip_cross_attn=False` re-enables cross-attention using the frozen (pre-trained) weights, which now operate on self-attention features adapted to the target domain.

#### Hyperparameters

LR ∈ [10^−6^, 10^−5^], weight decay 0, MMD loss only (ℒ= ℒ_MMD_), 200–1,024 NTC sets, cosine warmup 10%. Training is step-based (`max_steps` as the stopping criterion). A hyperparameter sweep over LR and total NTC sets (200, 500, or 1,024) is conducted per target domain (56 total configurations: 4 LRs × 2 set counts × 7 target datasets), with validation loss as the selection criterion.

#### Post-TTA inference

The adapted checkpoint is loaded for inference with `model_overrides:` {`skip_cross_attn: false`} to re-enable perturbation conditioning. DDP checkpoints (single `.pt` file) and FSDP checkpoints (sharded directory) are both supported transparently.

### A.6 Training Validation Metrics

The following metrics are computed on a held-out validation split during training to monitor optimization and detect collapse; they are *not* the final held-out test metrics reported in the main text. Training validation metrics operate on pseudobulk centroids aggregated within each predicted set of *S* = 64 cells at the subsampled context length (*G*^*′*^ ≤ 4,000), and fold-changes are computed relative to the matched control set. In contrast, cell-eval test metrics (Section 4.1.8) operate on full-context predictions (*G* ≈ 19k) from unseen datasets or contexts. Notably, training DE Pearson *r* flattens fold-changes across *all* perturbations and genes into one vector before computing correlation (a single global *r*), whereas cell-eval Pearson *δ* computes a per-perturbation correlation then averages. Similarly, 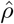 (delta norm ratio) is a training diagnostic for magnitude collapse (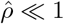) with no direct cell-eval counterpart.

#### Per-perturbation MAE

For each unique perturbation *p*, predictions are averaged over all cells with that perturbation to form a centroid, then compared to the true centroid:

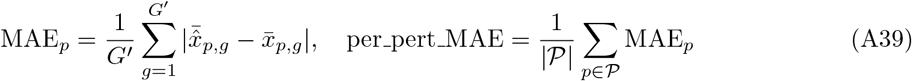

#### Centroid accuracy

For each predicted centroid 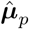, compute its L2 distance to the true centroid ***µ***_*p*_ versus all other true centroids within the same context. Centroid accuracy is the fraction of perturbations for which the nearest true centroid is the correct one (top-1 retrieval accuracy).

#### DE Pearson r

Fold-change vectors 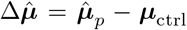 and Δ***µ*** = ***µ***_*p*_ − ***µ***_ctrl_ are flattened across all perturbations and genes, then Pearson correlation is computed. Measures whether the model captures the global pattern of differential expression.

#### Delta norm ratio ρ

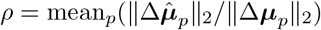. Values ≪ 1 indicate conservative collapse.

### A.7 Training Configuration

**Table A4.** Scaling experiment: architectures and final metrics (20 epochs).

| Config | $N$ (params) | Train loss | Val loss | DE-$r$ | $\hat{\rho}$ |
| --- | --- | --- | --- | --- | --- |
| 83M | 83,241,258 | 0.998 | 5.625 | 0.291 | 0.593 |
| 216M | 215,890,218 | 0.758 | 5.376 | 0.304 | 0.604 |
| 543M | 543,460,394 | 0.465 | 5.203 | 0.313 | 0.588 |
| 1.6B | 1,581,985,066 | 0.457 | 5.057 | **0.315** | 0.571 |
| 3.1B | 3,131,689,898 | 0.293 | **5.003** | 0.312 | 0.567 |

**Table A5.** Power-law fit parameters for *f* (*N*) = *a* · *N* ^−*α*^. Negative *α* indicates the metric *increases* with *N* .

| Metric | $\hat{a}$ | $\hat{\alpha}$ | $R^2$ |
| --- | --- | --- | --- |
| Train loss | 354.7 | +0.322 | 0.961 |
| Test loss | 10.09 | +0.033 | 0.971 |
| DE Pearson $r$ | 0.211 | -0.019 | 0.714 |
| MAE | 0.188 | +0.006 | 0.975 |

#### Batch size scaling

Phase 2 LR is derived from a base of 3 × 10^−5^ via square-root scaling [91]:

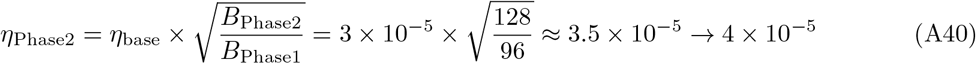

#### Phase 1 warmstart selection

Epochs 0, 6, and 9 of Phase 1 were evaluated on two held-out inference datasets (iPSC, ∼ 2,900 perturbations; APEL melanocyte differentiation, 34 curated test genes) via the cell-eval package. Epoch 9 won 8 of 12 cell-eval metrics across both datasets: Pearson *δ* improved from − 0.025 to 0.047 on iPSC and from 0.255 to 0.321 on APEL; DE direction match improved from 0.605 to 0.672 (iPSC) and 0.650 to 0.701 (APEL); DE recall improved from 0.662 to 0.747 (iPSC) and 0.578 to 0.626 (APEL). Despite diminishing returns in validation loss after epoch 6, the continued learning through the low-LR cosine tail (LR ≈ 3.4 × 10^−6^ at epoch 9) improved biological signal without harming generalization. The train/val gap (2.2× at epoch 9) reflected the high-effect data subset’s limited diversity; Phase 2’s fresh optimizer and 10× lower LR prevent amplification of any memorization [89].

#### scGPT pre-trained initialization

X-Cell initializes gene embeddings from a pre-trained scGPT model [8]. The vocabulary is mapped via ENSEMBL ID and symbol matching: pre-trained embeddings are transferred to matching genes in the new vocabulary, with unmapped genes retaining random initialization. The pre-trained transformer layers (12 layers, *d* = 512) are also loaded with `strict=False` to accommodate architectural differences. This warm initialization provides a strong starting point for gene-gene co-expression patterns.

### A.8 Distributed Training Details

#### FSDP2 configuration

Fully Sharded Data Parallel (FSDP2) [94] with per-layer wrapping on three module types: `ModernTransformerEncoderLayer`, `TransformerEncoderLayer`, and `CrossAttnBlock`. This achieves 74% of parameters in per-layer sharding units and 26% in the root (embeddings, decoder). `FULL_STATE_DICT` is used for checkpoints.

#### Activation checkpointing

Applied before FSDP preparation following the TorchTitan pattern [96]: activation checkpoint → `torch.compile` → FSDP sharding. This avoids dtype mismatch between FSDP2 and activation checkpoint wrappers.

#### Model FLOPs utilization

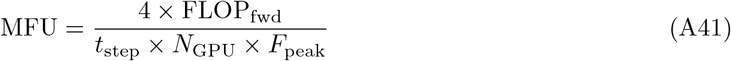

where the 4× accounts for 1× forward + 1× activation checkpoint recompute + 2× backward, and *F*_peak_ = 989 TFLOP/s (H200 bf16). X-Cell-Ultra achieves ∼41% MFU at 12.3 s/step on 64 H200 GPUs.

#### Training time estimation

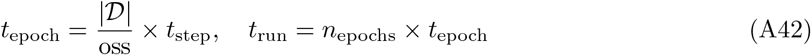

where |𝒟| is the number of cell sets in the dataset and oss is the optimizer step size (number of sets per gradient update). Per-layer FSDP wrapping reduced step time from ∼17 s to ∼12.3 s (28% improvement).

### A.9 Warmstart Vocabulary Remapping

When warmstarting a model trained on dataset 𝒟_*A*_ for fine-tuning on 𝒟_*B*_, gene vocabularies may differ. X-Cell constructs a remapping *π*: 𝒱_*B*_ → 𝒱_*A*_ via a multi-resolution lookup:

1. Match ENSEMBL IDs directly between feature sets.
2. Match gene symbols via GTF reference (symbol → ENSEMBL → feature index).
3. Unmapped genes receive the `<mask>` token index (shared embedding).

A 95% overlap threshold is enforced; warmstart fails if fewer than 95% of genes map successfully. The remapping tensor is injected into the collator so that `T gene` indices reference the pre-trained vocabulary at all times.

### A.10 Data Pipeline

#### A.10.1 Set Construction

The MMD dataloader constructs stratified sets of *S* cells for distribution-level training. For each (context, perturbation, batch) group, cell indices are permuted and chunked into sets of size *S*. Incomplete sets are padded by sampling with replacement. For each perturbed set, a matched control (NTC) set is sampled from the same context and batch, with fallback to the context-level NTC pool if batch-specific NTC cells are insufficient.

#### A.10.2 Preprocessing

Gene expression is preprocessed as: raw counts → CP10k normalization (*x*_*g*_ ← 10^4^ ·*x*_*g*_*/* ∑_*g*_ *x*_*g*_) → log(1 + *x*) → protein-coding gene filter via GRCh38-2024-A GTF reference. For pre-normalized datasets (Replogle-Nadig), CP10k and log1p are skipped. Gene subsampling to context length *G*^*′*^ ≤ 4,000 uses random selection with forced inclusion of the perturbed gene at position 1 (after CLS at position 0).

#### A.10.3 External Perturbation Embeddings

For each perturbation gene *p*, five pre-computed embedding vectors are loaded from reference files:

- **ESM-2** [38]: **z**_ESM_ ∈ ℝ^5120^, protein language model representation
- **STRING** [40, 41]: **z**_STRING_ ∈ ℝ^512^, protein-protein interaction network embedding
- **GenePT** [19]: **z**_GenePT_ ∈ ℝ^3072^, multi-modal gene representation from LLM
- **DepMap** [42]: **z**_DepMap_ ∈ ℝ^1150^, genetic dependency profiles across cancer cell lines
- **JUMP-Cell Painting** [43]: **z**_CP_ ∈ ℝ^259^, PCA of morphological phenotype features from U2OS cells

Each embedding is optionally LayerNorm-normalized before collation. Missing embeddings (gene not in source database) are zero-imputed and flagged via a boolean mask passed as `key padding mask` to cross-attention, so the model learns to ignore absent sources.

### A.11 Scaling Law Analysis

#### Experimental design

We trained five models of increasing capacity on Replogle-Nadig under identical hyperparameters: AdamW (*β*_1_ = 0.9, *β*_2_ = 0.999, *λ* = 0), peak LR *η* = 10^−4^, cosine schedule over *T* = 3,320 optimizer steps (20 epochs), set size *S* = 64, optimizer step size oss= 64, context length *G*^*′*^ = 4,000, on 8 nodes (64 H200 GPUs). Trainable parameter counts *N* were measured from the training logs:

Here DE-*r* is the Pearson correlation on differential expression fold-changes, with respect to control, and 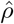 is the *δ*-norm ratio (predicted/true perturbation magnitude; ideal = 1.0, collapse *<* 0.2). Validation loss decreases monotonically with model size, and all models maintain 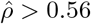 (no collapse).

#### Power-law fits

Following Kaplan et al. [58], we fit each metric as a power law *f* (*N*) = *a* · *N*^−*α*^ via nonlinear least squares on the five model sizes:

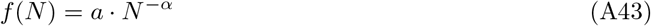

The train exponent 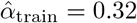 falls squarely within the range reported for language models (*α* ≈ 0.076– 0.34 [58, 59]), confirming that the X-Cell-Ultra architecture exhibits LLM-class scaling on single-cell perturbation data. Test loss scales with a shallower exponent (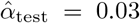, *R*^2^ = 0.97), reflecting diminishing returns as models approach the noise floor of the finite validation split. DE Pearson *r* has a *negative* exponent (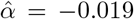), indicating that biological signal *improves* with scale, though the weaker fit (*R*^2^ = 0.71) and 1.0% plateau between 1.6B and 3.1B (*r* = 0.315 → 0.312) suggest saturation in this metric at current data scale. MAE decreases with a very shallow but clean power law (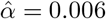, *R*^2^ = 0.98).

#### Data budget and compute-efficiency analysis

The MMD loss operates on *sets* of *S* cells, making the natural data unit a set rather than a token. Per optimizer step, the model processes oss = 64 sets × *S* = 64 cells/set × *G*^*′*^ = 4,000 genes/cell. Across 3,320 steps (20 epochs):

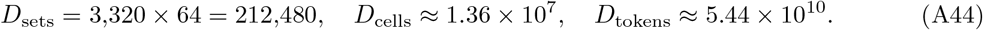

The per-model data-to-parameter ratios span nearly two orders of magnitude:

**Table A6.** Data-to-parameter ratios per model size. Chinchilla-optimal is 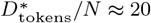 [59].

| Config | $D_{\text{tokens}}/N$ | $D_{\text{sets}}/N$ |
| --- | --- | --- |
| 83M | 307 | $2.5 \times 10^{-3}$ |
| 216M | 118 | $9.7 \times 10^{-4}$ |
| 543M | 47 | $3.9 \times 10^{-4}$ |
| 1.6B | 16 | $1.3 \times 10^{-4}$ |
| 3.1B | 8 | $6.7 \times 10^{-5}$ |

The smallest models are heavily overtrained in token terms (*D/N* = 654 for 83M), while the 3.1B model (*D/N* ≈ 17) approaches the Chinchilla-optimal *D*^∗^*/N* ≈ 20 [59]. The steep train exponent (*α* = 0.32) but shallow test exponent (*α* = 0.03) is consistent with this transition: smaller models have converged and are data-saturated, while the largest model still benefits from additional capacity. However, the set-level ratio *D*_sets_*/N* ≈ 7 × 10^−5^ for 3.1B remains orders of magnitude below unity, indicating that effective diversity is ultimately constrained by the ∼ 10,700 unique (perturbation, context) groups in the dataset rather than by total token count.

#### Validation loss and biological signal

Both train and test loss scale monotonically with model size (Table A4). The train–test gap remains roughly constant (Δ ≈ 4.6–4.7), indicating that 20 epochs provides sufficient training across all scales without catastrophic overfitting. DE Pearson *r* improves from 0.291 (83M) to a peak of 0.315 (1.6B), with 3.1B achieving *r* = 0.312 (within 1% of the 1.6B peak). This ∼ 8% improvement in fold-change correlation across a 37 × parameter range, combined with the near-Chinchilla *D/N* ratio for 3.1B, suggests that biological signal saturates before loss does—consistent with a regime where dataset diversity, not model capacity, is the binding constraint on downstream metrics.

## Appendix B Benchmarking implementation details

### B.1 Cell2Sentence-Scale Perturbation Prediction Benchmark

#### B.1.1 Cell2Sentence-Scale model

Cell2Sentence (C2S) converts single-cell gene expression profiles into natural language “cell sentences”. These sentences are space-separated gene names ordered by descending expression level, allowing autoregressive language models to predict cellular responses to perturbations (Levine et al., 2023; Rizvi et al., 2025). We benchmarked the larger-scale release of this model, “Cell2Sentence-Scale” and compared C2S-Scale-Gemma-2-2B and C2S-Scale-Gemma-2-27B, which are Gemma-2 language models pre-trained on over 57 million human and mouse cells from CellxGene and the Human Cell Atlas, against our model. Perturbation prediction is framed as a sequence-to-sequence translation task: given a control cell sentence and a perturbation label, the model generates the predicted perturbed cell sentence.

#### B.1.2 Data Preparation for cell sentences

For each benchmark dataset (Replogle-Nadig and Parse PBMC zero- and few-shot datasets), we split cells into non-overlapping train and test sets with perturbation-level holdout to support zero-shot evaluation claims. Gene identifiers were mapped from Ensembl IDs to HGNC gene symbols using the GRCh38-2024-A reference GTF. Expression matrices were pre-processed with log1p normalization (consistent with C2S pre-training conventions) and converted to Arrow-backed cell sentence datasets using the [cell2sentence] library (publicly available on GitHub).

##### Prompt Formatting and Control-Perturbed Cell Pairing

Each training example pairs a control cell with a perturbed cell. Control cells (labeled “non-targeting” for Replogle-Nadig or “PBS” for Parse) were pooled globally and paired with perturbed cells via uniform random sampling with replacement, following the approach described in the C2S perturbation prediction tutorial (Tutorial 10). No covariate matching (e.g., by batch, cell type, or other `.obs` metadata) was performed, consistent with the canonical C2S implementation. The prompt template was:

Given the following cell sentence of {num genes} expressed genes representing a cell’s basal state, predict the cell sentence after applying the perturbation: {perturbation name}. Control cell sentence: {control cell sentence}. Perturbed cell sentence:

The expected response was the perturbed cell’s top-K gene sentence followed by a period. We used the top 1,000 most highly expressed genes per cell (`top_k_genes=1000`), providing substantially more gene coverage than the 200-gene default in the C2S tutorial.

#### B.1.3 Training C2S-Scale models on new datasets

Fine-tuning was performed with full-parameter updates (no LoRA or adapter layers) using the HuggingFace Trainer with Accelerate and FSDP for multi-node distribution (on an H200 cluster, Oracle OCI). Training loss was computed only on response tokens (prompt tokens masked with label=- 100), following the `loss_on_response_only=True` convention in C2S. We selected the following **key hyper-parameters:**

- *context length:* 4,096 tokens total, split evenly between prompt and response (`response_ratio=0.5`). At approximately 4 tokens per gene name, this accommodates 500 of the 1,000 specified genes per cell sentence after truncation. Prompts were truncated independently from responses to guarantee that the model always has response tokens to learn from, preventing training collapse.
- *batch size:* 4 per device with 4 gradient accumulation steps across 16 GPUs (2 nodes of 8x H200), yielding an effective batch size of 256.
- *learning rate:* 1*e* − 6 for both the 2B and 27B model, with cosine decay schedule and 3% linear warmup
- *epochs:* 1 (single pass over the training set)
- *FSDP strategy:* `FULL_SHARD` with `TRANSFORMER_BASED_WRAP` on `Gemma2DecoderLayer`. Multi-node communication used NCCL over RDMA with Gloo for rendezvous.

Tokenized datasets were cached to shared and networked file-system storage and loaded with `keep_in_memory=True` to enable training from memory-mapped Arrow files on multi-node NFS. The HuggingFace model cache was set to a shared NFS directory to ensure all nodes access the same downloaded weights.

#### B.1.4 Distributed inference of C2S-Scale

Inference was performed using Accelerate’s `split_between_processes` for data-parallel distribution across GPUs. Each GPU loaded a full copy of the model in bfloat16 with Flash Attention 2 enabled (`device_map` set explicitly per rank; no FSDP). Predictions were generated using greedy decoding (`do_sample=False`’) with `max_new_tokens` set to half the total context length.

For zero-shot evaluation, we used the base pre-trained C2S model without fine-tuning. For fine-tuned evaluation, inference context length settings matched training (4,096 total, 0.5 response ratio). Predictions from each rank were saved as per-rank JSON files and aggregated by the main process with a polling-based timeout to handle potential rank failures gracefully and enable distributed inference for speed reduced as a fold-factor of the number of available GPUs.

#### B.1.5 Expression Reconstruction via Isotonic Regression

C2S generates ranked gene lists rather than expression values. To convert predicted gene orderings back to expression space for quantitative comparison, we fit an isotonic regression model mapping rank position to expression value, calibrated on control cells from the training set. Specifically:

1. For each control cell, expression values were sorted in descending order and the top-K values retained.
2. All (rank, expression) pairs across control cells were pooled and an `IsotonicRegression` model (implementation with `sklearn`, with `increasing=True`, and `out_of_bounds=‘clip’`) was fit on (rank, -expression) to learn the monotonically decreasing rank-abundance relationship.
3. For each predicted cell sentence, gene names were parsed in order, matched against the reference gene vocabulary, and assigned expression values from the isotonic model based on their predicted rank position. Genes not in the reference vocabulary were ignored; unmatched positions received NaN values.

This approach is more flexible than the linear rank-expression model (slope ∗ log_10_(rank + 1) + intercept) used in the original C2S reconstruction code, as isotonic regression captures the non-linear shape of empirical rank-abundance curves without imposing a parametric form.

#### B.1.6 Evaluation

Predicted and ground-truth expression matrices were compared using per-cell mean absolute error (MAE), computed only over genes present in both the prediction and reference. MAE was aggregated per perturbation and overall. Cells with zero matched genes (due to the model generating only out-of-vocabulary tokens) were flagged but included in reporting. All metrics were logged to Weights & Biases.

---

**Supplementary Figure 1.**
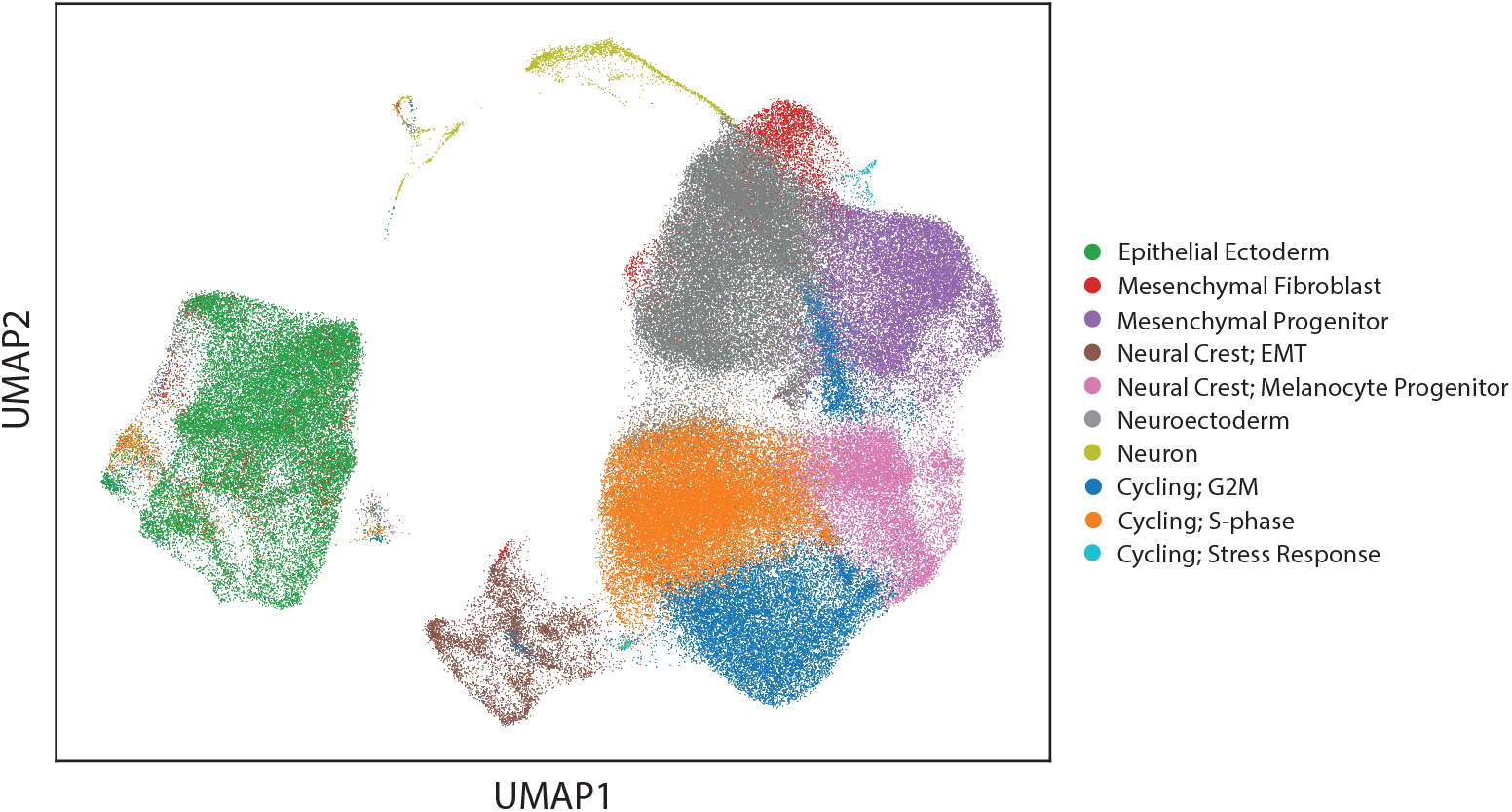
UMAP of iPSC Multi-Diff screen colored by cell type. Data from one GEM is shown. For details on cell type annotation, see Section 4.2.12.

**Supplementary Figure 2.**
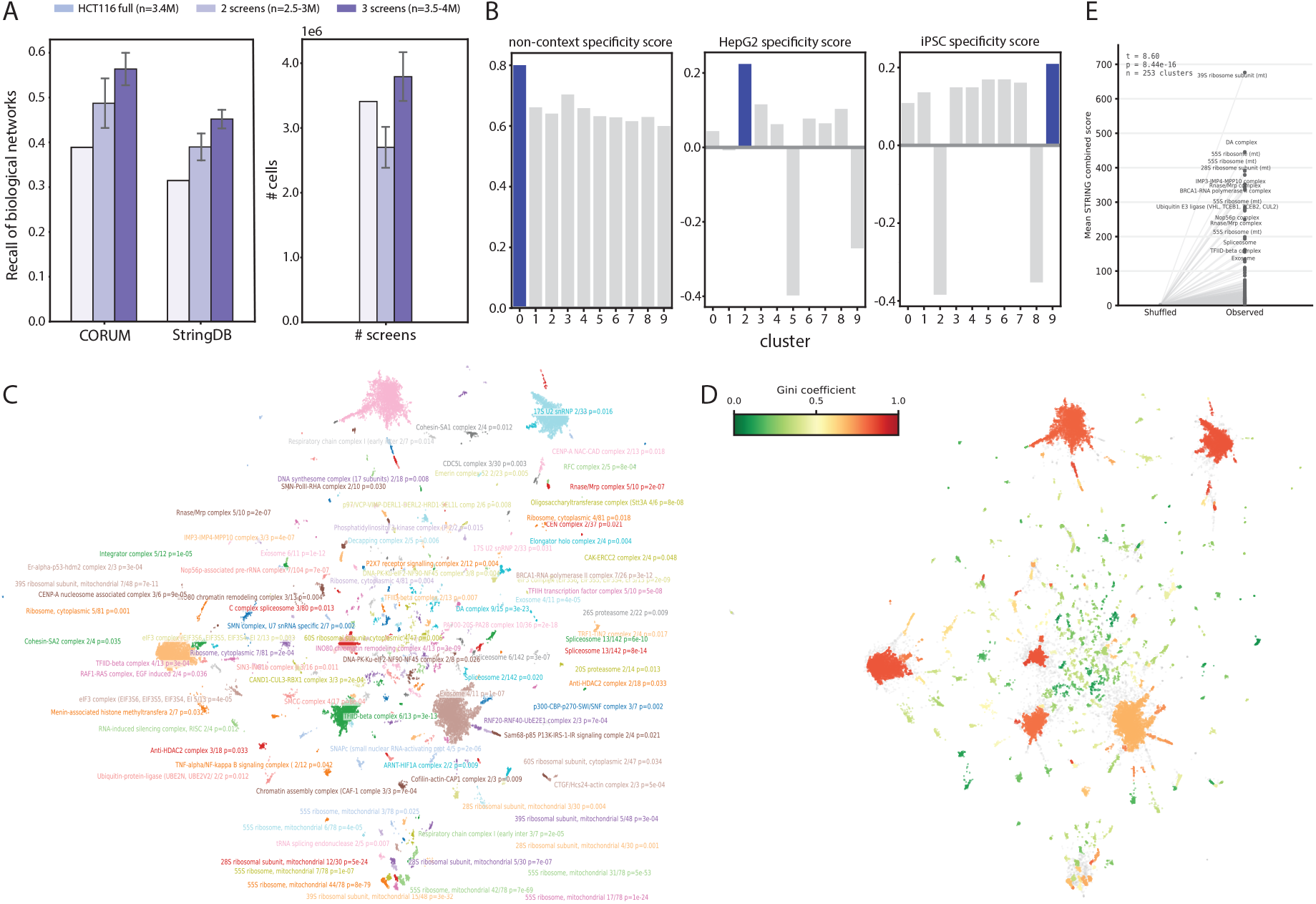
X-Atlas/Pisces enables identification of context-dependent and context-independent perturbations. **(A)** Recall of annotated gene pairs from CORUM and STRING, calculated from (1) the full HCT116 screen, (2) downsampled two screen combinations containing HCT116, and (3) downsampled three screen combinations containing HCT116 (left). Bargraph of number of cells in each screen combination (right). **(B)** Bargraph of non-context specificity scores (mean F1 score across screens) (left), HepG2 specificity scores (HepG2 F1 score - mean F1 score from other screens) (middle), and iPSC specificity scores (iPSC F1 score - mean of F1 scores from other screens) (right) across clusters. **(C)** UMAP of significant perturbations colored by screen. Clusters were automatically annotated with their corresponding CORUM protein complex based on gene set overlap. Adjusted p-values were calculated using Fisher’s exact test with Benjamini-Hochberg correction. **(D)** UMAP of significant perturbations colored by Gini coefficient across screens. **(E)** Mean STRING scores within clusters (observed) compared to that of 200 random permutations of perturbation labels (shuffled). Paired Student t-test was used to determine significance (t-test statistic=8.60, *p*-value = 8.44*e*^−16^).

**Supplementary Figure 3.**
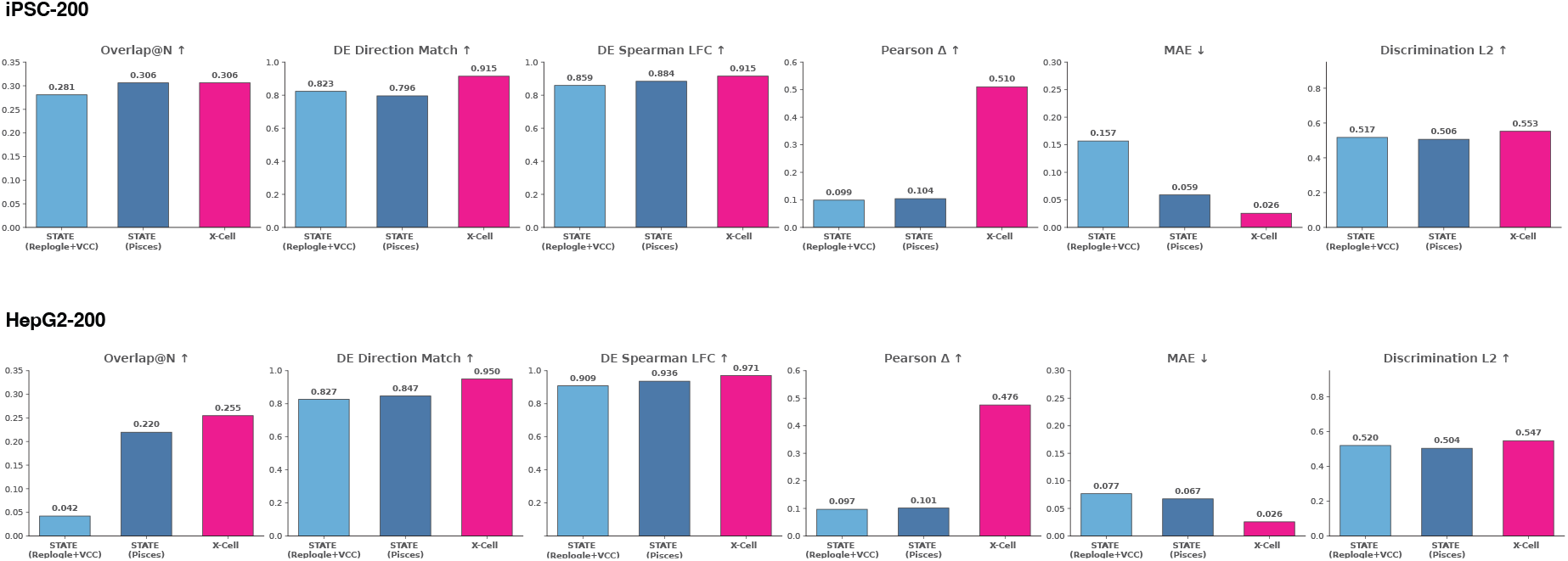
Performance comparison between STATE models trained on different datasets and X-Cell. For a fair comparison, we trained STATE (Pisces) model on a dataset set matching that used to train the X-Cell model, and STATE replogle on all publicly available datasets. We evaluate performance using six metrics across iPSC and HepG2 cell lines.

**Supplementary Figure 4.**
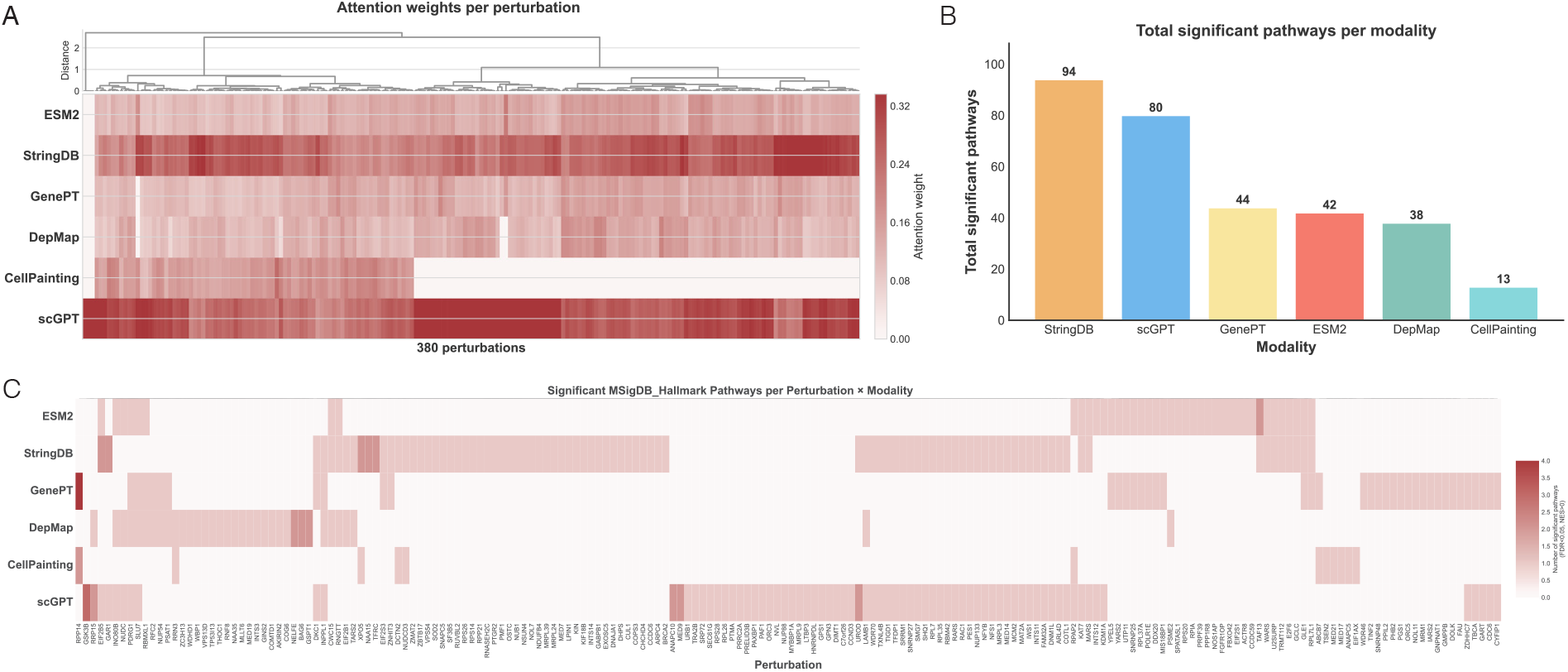
Characterization of Prior Knowledge Sources. (A) A heatmap of attention weights, showing allocation across 380 perturbations of *Replogle-Nadig* HepG2 test set (columns) and six knowledge sources (rows). Perturbations are hierarchically clustered using Ward linkage to group similar attention profiles. Color intensity represents attention weight, with the scale capped at the 95th percentile to minimize the effect of outliers. **(B)** A bar chart summarizing the total count of significant MSigDB Hallmark pathways captured by each knowledge source. Gene attention scores were used as the ranking metric for GSEA, with significant pathways defined by an FDR *<* 0.05 and a positive Normalized Enrichment Score (NES). **(C)** A detailed heatmap that breaks down the GSEA results from (b). It displays the number of significant pathways for each individual perturbation, illustrating how the biological relevance of signals from each knowledge source varies across different conditions.

**Supplementary Figure 5.**
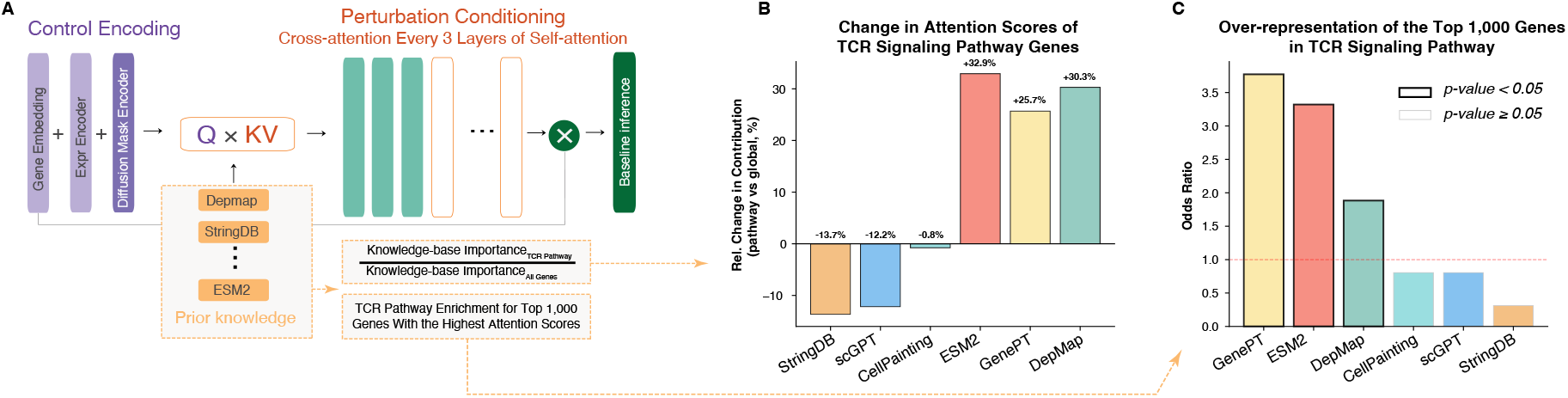
External knowledge bases provide context-specific information to X-Cell. **(A)** Schematic of the X-Cell architecture, which employs cross-attention between gene embeddings (initialized via scGPT) and external knowledge bases including DepMap, STRING, CellPainting, ESM-2, and GenePT. To evaluate the contribution of relevant knowledge from these sources, we analyzed cross-attention scores during inference of the *CD3E* perturbation (a known T cell inactivating perturbation) among activated Jurkat cells. We compared the distribution of attention scores for genes within the KEGG TCR Signaling pathway (hsa04660) against the global distribution of all genes. An upward shift in scores indicates the increased importance of these knowledge sources for this specific pathway. **(B)** Bar plots illustrating the increase in attention scores for ESM-2, GenePT, and DepMap embeddings within the KEGG TCR signaling pathway genes. **(C)** Over-representation of the top 1,000 genes with the highest attention scores for each modality within the KEGG TCR signaling pathway. Bars represent the Odds Ratio (OR) from a one-sided Fisher’s exact test; black outlines denote statistical significance (*p <* 0.05).

**Supplementary Figure 6.**
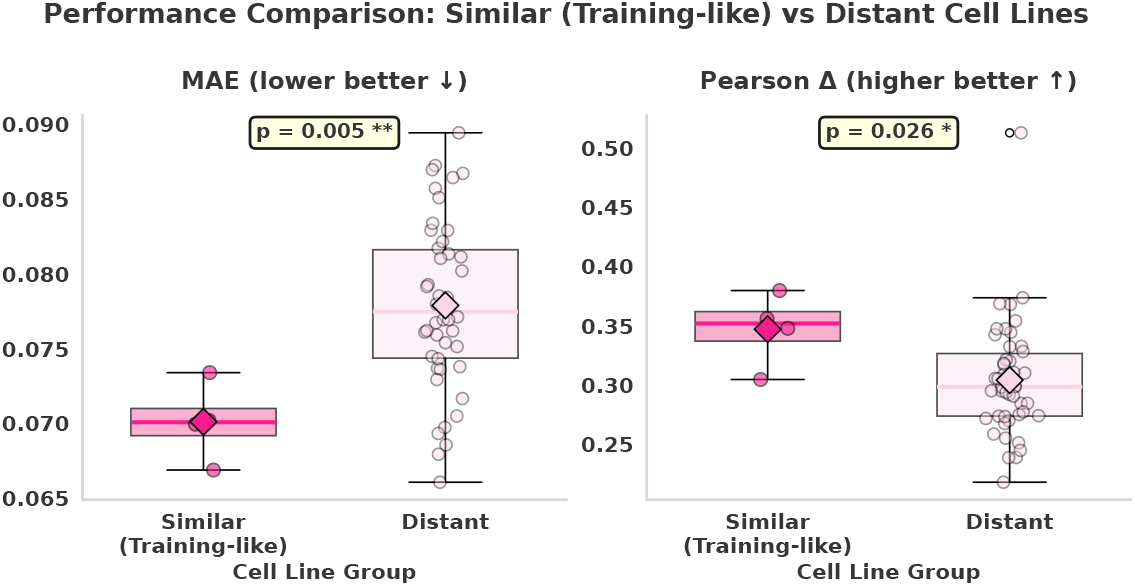
Comparison of zero-shot model performance on Tahoe-100M dataset between four cell lines most similar to training (pink; HCT15, RKO, SW480, and HepG2/C3A) and distant cell lines (light pink). Box plots show distributions for each group, with mean values indicated. Statistical significance is assessed using the Mann-Whitney U test.

**Supplementary Figure 7.**
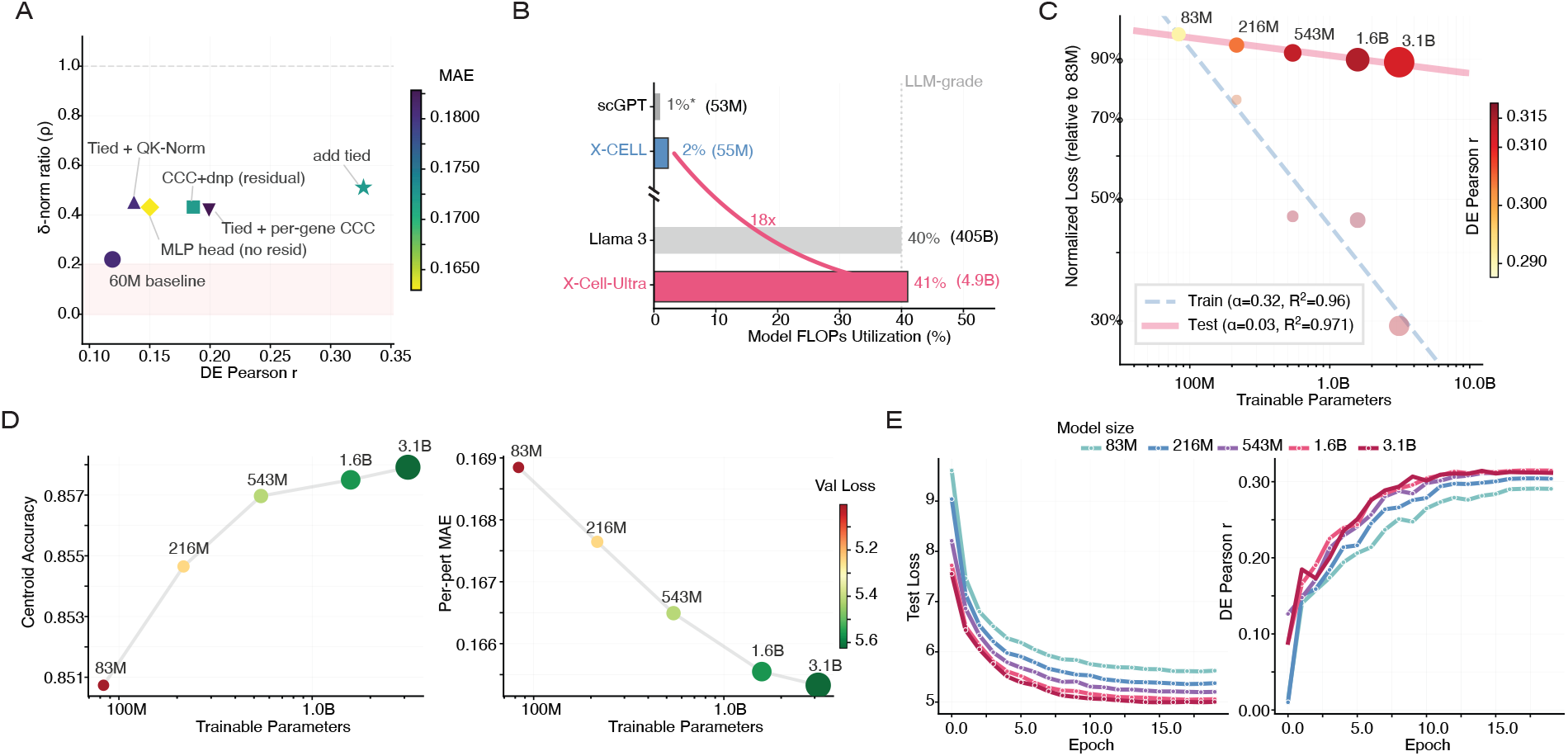
Architecture ablations, training efficiency, and scaling analysis for X-Cell-Ultra. **(A)** Architecture and loss ablation on the 4.9B model (Replogle-Nadig, 6 epochs). Scatter shows *δ*-norm ratio 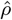 vs. DE Pearson *r* for six configurations; color = MAE. The red shaded region 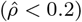 indicates conservative collapse (predicting small effects). Replacing scale-invariant losses with CCC and removing the hidden residual shortcut progressively recovered both magnitude 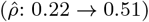 and direction (DE-*r*: 0.05 → 0.33); see Table A1. **(B)** Model FLOPs Utilization (MFU) comparison. X-Cell-Ultra achieves 41% MFU on 64 H200 GPUs, comparable to Llama 3 (reported 38–43% [95]) and ∼ 20 × the 55M X-Cell (2%). **(C)** Normalized loss scaling (relative to 83M model). Both train (dashed, *α* = 0.32, *R*^2^ = 0.96) and test (solid, *α* = 0.03, *R*^2^ = 0.97) follow power laws; color = DE Pearson *r*. **(D)** Individual validation metrics vs. trainable parameters: centroid accuracy (left) and per-perturbation MAE (right), colored by validation loss. **(E)** Training dynamics over 20 epochs: test loss (left) and DE Pearson *r* (right) for all five model sizes.

**Table B7.** Benchmark performance across evaluation datasets in Section 2.3. Best values per metric are shown in bold. cell-eval metrirc are defined in Section 4.1.8.

| Metric | C2S-2B | STATE | scGPT | X-Cell |
| --- | --- | --- | --- | --- |
| **iPSC-200** |  |  |  |  |
| Overlap@N ↑ | 0.0453 | 0.3062 | 0.1676 | **0.3064** |
| DE Direction Match ↑ | 0.7128 | 0.7958 | 0.8018 | **0.9149** |
| DE Spearman LFC ↑ | 0.8014 | 0.8840 | 0.8635 | **0.9153** |
| Pearson $\Delta$ ↑ | -0.0327 | 0.1042 | 0.0160 | **0.5104** |
| MAE ↓ | 0.0654 | 0.0590 | 0.0304 | **0.0257** |
| Centroid Accuracy ↑ | 0.5045 | 0.5062 | 0.5352 | **0.5525** |
| **HepG2-200** |  |  |  |  |
| Overlap@N ↑ | 0.0519 | 0.2196 | 0.0893 | **0.2548** |
| DE Direction Match ↑ | 0.7971 | 0.8470 | 0.8588 | **0.9502** |
| DE Spearman LFC ↑ | 0.8782 | 0.9363 | 0.9650 | **0.9710** |
| Pearson $\Delta$ ↑ | -0.0061 | 0.1014 | 0.0491 | **0.4756** |
| MAE ↓ | 0.0638 | 0.0673 | 0.0299 | **0.0256** |
| Centroid Accuracy ↑ | 0.4940 | 0.5038 | 0.5244 | **0.5473** |
| **Replogle-Nadig** |  |  |  |  |
| Overlap@N ↑ | 0.1432 | 0.2600 | 0.2013 | **0.3053** |
| DE Direction Match ↑ | 0.6007 | 0.7564 | 0.6867 | **0.8762** |
| DE Spearman LFC ↑ | 0.3369 | 0.5798 | 0.4298 | **0.7745** |
| Pearson $\Delta$ ↑ | 0.0538 | 0.4563 | 0.1884 | **0.5137** |
| MAE ↓ | 0.2845 | 0.0864 | 0.0804 | **0.0682** |
| Centroid Accuracy ↑ | 0.5023 | 0.5767 | 0.6786 | **0.7876** |
| **Parse-1M** |  |  |  |  |
| Overlap@N ↑ | 0.1476 | **0.3044** | 0.2090 | 0.2500 |
| DE Direction Match ↑ | 0.9114 | **0.9478** | 0.9435 | 0.9459 |
| DE Spearman LFC ↑ | 0.8298 | 0.9227 | 0.9283 | **0.9414** |
| Pearson $\Delta$ ↑ | 0.3625 | 0.5360 | 0.4459 | **0.5858** |
| MAE ↓ | 0.1492 | 0.1036 | 0.0941 | **0.0850** |
| Centroid Accuracy ↑ | 0.7547 | 0.6995 | 0.6828 | **0.9144** |

## Footnotes

1 https://zenodo.org/records/10833191/

2 https://stringdb-downloads.org/download/protein.network.embeddings.v12.0.h5

3 https://plus.figshare.com/articles/dataset/DepMap_24Q4_Public/27993248

4 https://www.biorxiv.org/content/10.1101/2023.03.23.534023v2

5 https://plus.figshare.com/articles/dataset/Mapping_information-rich_genotype-phenotype_landscapes_with_genome-scale_Perturb-seq_Replogle_et_al_2022_processed_Perturb-seq_datasets/20029387

6 https://www.ncbi.nlm.nih.gov/geo/query/acc.cgi?acc=GSE264667

7 https://virtualcellchallenge.org/datasets

8 https://huggingface.co/arcinstitute/SE-600M/tree/main

9 https://huggingface.co/arcinstitute/ST-SE-Replogle/tree/main/zeroshot

10 https://colab.research.google.com/drive/1Ih-KtTEsPqDQnjTh6etVv_f-gRAA86ZN

11 https://huggingface.co/arcinstitute/ST-SE-Parse

12 https://github.com/vandijklab/cell2sentence/blob/master/tutorials/c2s_tutorial_10_perturbation_response_prediction.ipynb

13 https://github.com/bowang-lab/scGPT/blob/main/tutorials/Tutorial_Perturbation.ipynb

14 https://huggingface.co/datasets/arcinstitute/Replogle-Nadig-Preprint/resolve/main/replogle.h5ad

15 https://colab.research.google.com/drive/1Ih-KtTEsPqDQnjTh6etVv_f-gRAA86ZN

16 https://huggingface.co/datasets/arcinstitute/State-Parse-Filtered/resolve/main/parse_concat_full.h5ad

17 https://huggingface.co/arcinstitute/ST-SE-Parse/blob/main/fewshot/split_3.toml

18 https://huggingface.co/datasets/tahoebio/Tahoe-100M/viewer/drug_metadata

